# WTAP-mediated epitranscriptomic program in alveolar macrophages confers prolonged protection against postinfluenza bacterial pneumonia

**DOI:** 10.64898/2026.05.23.727154

**Authors:** Yong Ge, Xinyi Hu, Zhen Li, Youxiang Cheng, Rong Chen, Huan Wu, Zekai Qian, Wenwen Song, Jingrong Huang, Yilin Zou, Nan Qi, Anlong Xu, Shaochun Yuan

## Abstract

Secondary bacterial infections remain an intractable problem of seasonal influenza, primarily due to the depletion and defunctionalization of alveolar macrophages (AMs). Here, we described that *N^6^*-methyladenosine (m^6^A), the most prevalent RNA modification in eukaryotes, critically orchestrates trained immunity (TRIM) establishment in AMs. Influenza A virus (IAV)-trained AMs maintained low Wilms tumor 1-associated protein (WTAP) expression and reduced global m^6^A deposition for over two months. Mechanistically, this IAV-induced m^6^A decrease promotes TRIM by enhancing the RNA stability of phagocytic and metabolic genes, thereby boosting antibacterial function. Mimicking this m^6^A reduction pharmacologically or genetically recapitulates the IAV-trained TRIM phenotype, which improves phagocytosis and protects mice from secondary infection. Clinically, elevated WTAP in AMs correlates with impaired phagocytosis and disease severity in COVID-19 and COPD. These findings for the first time unveil how respiratory virus infection shapes AM TRIM via epitranscriptomic reprogramming, and offer prospective strategies for the prevention and treatment of post-viral bacterial complications.

## Introduction

Lower respiratory infections (LRIs) kill more than 2 million individuals each year, making it the fourth leading cause of deaths in the world (1). Influenza virus is a major contributor to this burden, driving recurrent seasonal epidemics (2). Severe influenza viral disease is caused by direct cytopathic effects, aggravated host immune response to high viral loads, and secondary bacterial pneumonia resulting from acute injury (3, 4). Although most influenza patients recover within 1-2 weeks after infection, the prolonged impact of this ubiquitous infection on lung immunity has gradually aroused clinician’s attention. For example, influenza can shape pulmonary immunity by altering alveolar macrophages (AM) function in ways that contribute to chronic lung diseases (5). Recent studies further indicate that influenza-trained AMs can develop innate immune memory, influencing outcomes of secondary lung diseases, such as bacterial pneumonia or cancer (6, 7). Thus, intervention strategies built upon the functional remodeling of AMs are particularly important for achieving protective immunity against secondary bacterial pneumonia.

As resident sentinels of the alveoli, AMs are the first innate immune cells to encounter inhaled pathogens (8). They play a central role in maintaining lung homeostasis by phagocytizing inhaled pathogens and cellular debris at the air-tissue interface of lung alveoli, thereby restraining excessive inflammation (9). In the steady state, AM functions are regulated in part by nuclear receptor peroxisome proliferator-activated receptor γ (PPAR-γ). PPAR-γ signaling in AMs upregulates phagocytosis through both CD36 and Fcγ receptors (FcγR) while suppresses pro-inflammatory activities by inhibiting the transcription of a subset of inflammatory genes (10). Once pathogen containment fails, AMs can initiate a well-orchestrated inflammatory response, and promptly resolve inflammation to restore homeostasis in the aftermath of infection (11). These properties position AMs as an attractive candidate for novel therapies against pulmonary infection and related diseases. For example, influenza infection impairs AM crawling and increases secondary bacterial co-infection (12, 13), so enhancing AM chemotaxis toward bacteria may avoid superfluous neutrophil recruitment and inappropriate inflammatory injury. Furthermore, given the anti-inflammatory and pro-repair properties of AMs, adoptive transfer of AMs derived from pluripotent stem cells (PSCs) may also be a successful strategy in combating pulmonary infections (14, 15). Yet, AM-based cell therapies are still in their infancy, underscoring the need to better understand the functional reprogramming of AMs in disease contexts.

Recently, long-term functional remodeling of AMs following influenza infection or vaccination (16), termed trained immunity (TRIM), has gained attention for its role in modulating secondary bacterial infections and lung immune homeostasis (17). TRIM reflects the sustained hyper-responsiveness of innate immune cells, which provides a mechanism for the rapid transcriptional activation upon re-challenge (18). Although TRIM can be induced by microorganisms and microbial products (*Candida* a*lbicans*, β-glucan, and BCG vaccine) or self-derived molecules (uric acid, oxLDL, and cytokines), these endogenous or exogenous compounds can lead to distinct functional adaptation of innate immune cells (19–22). The trained phenotype is established via characteristic epigenetic imprints that are fueled and regulated by coordinated metabolic rewiring. Metabolic rewiring provides necessary energy and metabolic intermediates (23, 24), while concurrent epigenetic reprogramming including H3K27ac and H3K4me3 increases chromatin accessibility to potentiate a tailored gene transcription (25). The great potential of TRIM for clinical application lies in its dual capacity to promote cross-protection and resolve non-infectious inflammation (26). Insights into the specific molecular mechanism responsible for AM TRIM could therefore facilitate the development of novel therapeutic strategies that target a broad range of respiratory pathogens.

Although the transcriptional control of AM TRIM mediated by epigenetic and metabolic interweaving has been studied extensively, the importance of post-transcriptional regulation of these processes is less well defined. *N*^6^-methyladenosine (m^6^A), the most abundant internal chemical modification on eukaryotic mRNA, has emerged as a key post-transcriptional regulator of immune cell fate and function (27). m^6^A is installed on mRNAs by a methyltransferase complex called ‘writer’ (e.g., METTL3/METTL14/WTAP), and removed by a demethylase namely ‘eraser’ (FTO and ALKBH5). The resulting modification is then interpreted by distinct ‘readers’, such as YTH and IGF2BP proteins, which influence RNA splicing, stability, export, and translation to fine-tune gene expression at the epitranscriptomic level (28). There is still no direct evidence linking m^6^A to TRIM establishment, but m^6^A-mediated regulation of TNF mRNA stability in macrophages supports prolonged inflammatory remodeling during chronic inflammation (29). Moreover, the m^6^A methyltransferase inhibitor STC-15 can induce durable anti-tumor immunity, possibly via immune memory formation (30, 31). Thus, whether and how RNA epitranscriptomic mechanisms contribute to TRIM in AMs represents a significant knowledge gap.

In this study, we investigated whether RNA epitranscriptomic modifications underlie the long-term imprint in AMs after influenza infection. We found that influenza-experienced mice harbor an AM population with downregulated WTAP expression and globally reduced m^6^A deposition, a state that persists for at least two months. Influenza-trained or WTAP-deficient AMs entered a pronounced TRIM state, characterized by enhanced phagocytic and hydrolytic capacity through integrated epitranscriptomic and metabolic rewiring. These reprogrammed AMs promoted non-inflammatory clearance and limited inflammatory lung injury during secondary bacterial challenge. Exploring the mechanism of AM TRIM, we first identified WTAP-dependent m^6^A as a key regulator and further demonstrated that reducing m^6^A abundance significantly enhances the trained phenotype. Abolishment of the m^6^A modification in many phagocytic and metabolic genes, such as *Pparg*, *Hk2*, *Fcgr3* and *Fcgr4*, could enhance their mRNA stability and protein production, ultimately boosting cellular phagocytic and metabolic capacity. Collectively, our study identifies an RNA epitranscriptomic mechanism driving TRIM in AMs, offering new theoretical foundations and potential therapeutic targets for preventing and treating secondary pulmonary infections.

## Results

### IAV-trained AMs are characterized by a high phagocytic capacity

Exposure to respiratory viruses like influenza A (IAV) and SARS-CoV-2 can establish TRIM in AMs, however, the mechanistic basis driving this process remains poorly defined. To address this, we first analyzed a published single-cell RNA sequencing (scRNA-seq) dataset [accession nos. GSE208294] of bronchoalveolar lavage (BAL) CD11c^+^ cells from mice at baseline, day 7, day 14, day 21 and day 35 post-IAV administration (32). Unsupervised clustering over all time points distributed cells into 4 transcriptionally distinct clusters of CD11c^+^ cells, identified by signature genes (Figure 1A). Of note, tissue-resident AMs (TR_AMs) were widely depleted on day 7 after IAV infection and gradually returned between day 14 to day 35, while the number of monocyte-derived AMs (Mo_AMs) showed an opposite trend (Supplementary Figure 1A), indicating that the dominant cell population in the alveoli is TR_AMs during the recovery stage of IAV infection. So, we next examined the long-term effects of IAV on TR_AMs. Using the Leiden algorithm at a resolution of 0.2 (a setting that achieved stable, biologically interpretable partitions while avoiding over-clustering), TR_AMs were further divided into 9 transcriptionally distinct subclusters (Figure 1B). Notably, the number and proportion of subcluster 0 were depleted at day 7 p.i., and gradually returned and markedly increased between day 14 to day 35 p.i. (Figure 1, C and D), indicating that subcluster 0 was the predominant TR_AMs in the lungs of mice in the recovery phase of infection. Importantly, in addition to common infection and immune-related pathways, gene clusters associated with phagocytosis, cell adhesion, antigen processing and presentation were also highly expressed in subcluster 0 TR_AMs (Figure 1, E and F, and Supplementary Figure 1, B and C). To assess the long-term effects of a transient viral infection on AM phagocytic function, we next analyzed the publicly available RNA-seq dataset [accession nos. GSE208294] of AMs from mice at baseline and day 28 post-IAV administration (7). The results showed that many genes related to phagocytosis in the AMs after IAV training showed significant upregulation (Supplementary Figure 1D). Earlier report has identified that the memory phase of bacterial infection in AMs is characterized by heightened phagocytic activity and increased scavenger receptor expression, which facilitates inflammation resolution by promoting efficient clearance of cellular debris (33). Our data analysis indicated that respiratory virus-trained AMs may also display enhanced phagocytic function.

**Figure 1.**
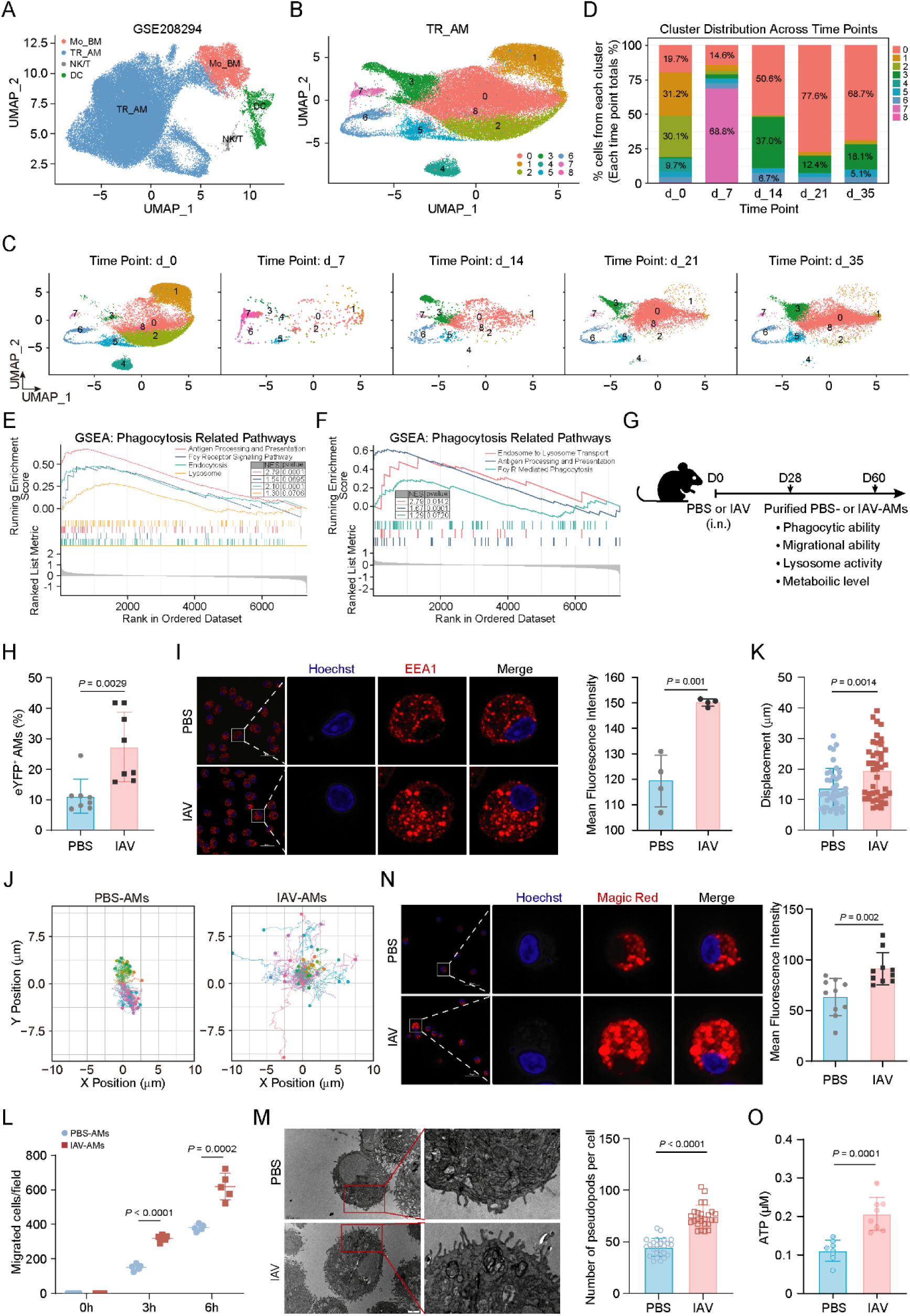
IAV-trained AMs are characterized by a high phagocytic capacity. (**A**) Uniform manifold approximation and projection (UMAP) map presentation of major cell types in BALFs from all mice; each dot represents a single cell. (**B**) UMAP displaying 9 TR_AM cell clusters (pooled data from all time points). (**C**) UMAP displaying 9 TR_AM cell clusters, and colors represent different cell populations at the indicated time points p.i. (**D**) Stacked bar plots showing the percentage of each cell clusters during the infection course. (**E** and **F**) GSEA enrichment of phagocytosis-related genes in cluster 0 from IAV-infected mice on day 21 (E) or day 35 (F) after infection versus those from uninfected mice. (**G**) Experimental schema of i.n. IAV infection in mice and subsequent phenotypic detections. (**H**) Flow cytometry quantifying the phagocytosis of PBS-and IAV-AMs on eYFP-*E. coli*. n = 8 mice per group. (**I**) Representative imaging of EEA1 immunofluorescence staining in PBS- and IAV-AMs (left) and quantification of EEA1 intensity (right). Scale bar, 25 μm. (**J**) Spider plot of randomly chosen PBS- and IAV-AMs, and colors represent individual AMs. (**K**) Quantification of net displacement (micrometers) of PBS- and IAV-AMs during 1 h imaging session. (**L**) Quantification of the *in vitro* chemotaxis assay of PBS- and IAV-AMs. n = 5 mice per group. (**M**) Representative TEM micrograph of pseudopods in AMs from control or IAV-infected mice (left) and quantification of pseudopod numbers (right). Scale bar, 2 μm. (**N**) Representative imaging of Magic Red dye in PBS- and IAV-AMs (left) and quantification of Magic Red intensity (right). Scale bar, 25 μm. (**O**) Determination of ATP concentration in PBS- and IAV-AMs. Data are presented as the mean ± s.d. in H, I, K and L-O with individual measurements overlaid as dots, and statistical analysis was performed in performed in H, I, K and M-O using a two-tailed Student’s *t*-test or performed in L using Multiple Comparisons Following Two-Way ANOVA. Data in I, M and N are representative of three independent biological experiments.

To assess the long-term effects of a transient viral infection on AM phagocytic function, we isolated and purified AMs (Supplementary Figure 2, A and B) from wild type C57BL/6J mice that were intranasally (i.n.) infected with IAV or endotoxin-free PBS for 28 days and analyzed their activity (Figure 1G). The phagocytic capacity of AM cells was first measured by incubating with *E. coli* expressing yellow fluorescent protein (eYFP-*E*. *coli*) followed by flow cytometry (Supplementary Figure 2C). The results revealed that phagocytosis of eYFP-*E*. *coli* was significantly higher in IAV-trained AMs than that in PBS controls (Figure 1H and Supplementary Figure 3A). Consistent with this, immunofluorescence staining of the early endosomal markers EEA1 (Early Endosomal Antigen 1) revealed that IAV-trained AMs form more early phagosomes after *P. aeruginosa* infection (Figure 1I). We used time-lapse microscopy to examine the consequences of IVA training on AM migration. As expected, IAV-trained AMs had a significant increase in displacement compared with PBS controls (Figure 1, J and K, and Supplementary Figure 3, B and C). AMs crawl in and between alveoli to phagocytose inhaled bacterial pathogens with high efficiency (12), so we next performed *in vitro* migration assays (Supplementary Figure 3D). Transwell assays also showed increased migratory events by IAV-trained AMs after coincubation with *P. aeruginosa* (Figure 1L and Supplementary Figure 3E). Macrophages form pseudopods to facilitate the ingestion and clearance of bacteria (34), our transmission electron microscopy (TEM) results also showed that IAV-trained AMs have more pseudopodia than PBS controls (Figure 1M). To further measure the persistence of this phagocytic function, we isolated IAV-trained AMs from mice at 60 days post-infection (Figure 1G). As we expected, such a high migratory and phagocytic phenotype was consistently observed in trained AMs through day 60 after IAV infection (Supplementary Figure 3, F to H). These data suggested that IAV-trained AMs have a long-lasting and stronger ability to phagocytose bacteria. Given that macrophages digest phagocytosed bacteria via phagolysosome, we further measured the abundance and activity of lysosomes in PBS- or IAV-trained AMs. Using cell-permeant Magic Red to measure the activity of cathepsin B, a lysosomal marker enzyme, we found that IAV-trained AMs had higher Magic Red fluorescence (Figure 1N), indicating enhanced hydrolytic function of lysosomes. In support of enhanced phagocytic and hydrolytic effects, increased energy metabolism is required in AMs (35). Therefore, we wanted to explore if there is also an adaptive change in the metabolism of IAV-trained AMs. As expected, ATP levels measured with luciferin assays revealed that IAV training resulted in higher cellular ATP content (Figure 1O and Supplementary Figure 3I). These data thus indicated that IAV-trained AMs exert antibacterial functions via enhanced phagocytosis and metabolism.

We next sought to assess whether IAV infection induces protection against heterologous bacterial infection in mice, as shown in previously studies (7, 16). To do this, mice were challenged intratracheally (i.t.) with a sublethal dose of *P. aeruginosa*, the leading cause of infections in immunocompromised individuals and in healthcare settings (36), on day 28 after IAV infection (Supplementary Figure 4A). As we expected, IAV-infected mice showed decreased body weight loss (Supplementary Figure 4B) and decreased BAL fluid (BALF) bleeding (Supplementary Figure 4, C-E) and reduced lung bacterial burdens (Supplementary Figure 4F) compared to PBS mice. The corresponding flow cytometry demonstrated reduced neutrophil infiltration (Supplementary Figure 4, G and H), which was corroborated with reduced inflammation and immunopathology (Supplementary Figure 4, I and J) in lung tissues from mice infected with IAV. These data indicated that mice with IAV-trained AMs exhibit enhanced phagocytic and hydrolytic function, conferring protection against bacterial pneumonia.

### WTAP is downregulated in AMs during IAV-induced trained immunity

Post-influenza mice acquire prolonged antibacterial resistance, a phenotype we confirmed and associated with stronger phagocytic and hydrolytic activity in AMs. We next investigated whether RNA epitranscriptomic modifications contribute to this long-term imprint, in addition to the documented epigenetic and metabolic rewiring after influenza infection. To answer this question, we first tested the abundance of m^6^A modification in PBS- and IAV-trained AMs. Liquid chromatography–mass spectrometry (LC–MS/MS) assays revealed a significant reduction in the overall m^6^A abundance in AMs from IAV-infected mice at day 28 post-infection (Figure 2A and Supplementary Figure 5A). In addition, decreases in m^6^A abundance were also confirmed by immunofluorescence staining (Figure 2, B-D). To further confirm this phenomenon, we analyzed the expression of m^6^A writers, erasers and readers in the TR_AMs from IAV-infected mice using aforementioned scRNA-seq dataset [accession nos. GSE208294]. The results revealed dynamic changes in the expression of most m^6^A regulators in TR_AMs during the course of IAV infection (Figure 2E). Notably, WTAP, the key regulatory subunit of the writer complex (37), was upregulated during acute IAV infection and downregulated in the recovery phase, showing a pattern that mirrored the overall m^6^A changes (Figure 2E). As we previously identified WTAP is a pro-inflammatory driver(38), its acute-phase induction likely contributes to the establishment of lung inflammation. On the contrary, a persistent significant reduction in WTAP in AMs at day 21 or even until day 35 post-IAV infection, implying that WTAP may also be involved in the functional remodeling of AMs during recovery from influenza infection. Subsequently, we further purified AMs from wild type C57BL/6J mice that were intranasally (i.n.) infected with IAV or endotoxin-free PBS for 28 days and performed RT-qPCR analyses to measure the expression of WTAP. In line with the above results, *Wtap* mRNA expression was downregulated in IAV-trained AMs (Figure 2F), and the protein abundance showed the same trend (Figure 2, G and H, and Supplementary Figure 5B). Thereafter paraffin slides of mouse lung tissue were stained with fluorescent antibodies to identify the WTAP expression in IAV-trained AMs, and the same trend was obtained (Figure 2, I-K). These data suggested that IAV-trained AMs are characterized by the decreased m^6^A abundance caused by the down-regulated WTAP. We also evaluated the adaptive changes of WTAP/m^6^A in AMs from IAV-infected mice at day 60 post-infection (Supplementary Figure 5A), and as expected, both reduced WTAP expression and decreased m^6^A modification remained evident on day 60 after IAV infection (Figure 2, L-N, and Supplementary Figure 5, C-F), indicating that a transient IAV infection induces a long-lasting innate immune memory state in AMs characterized by sustained downregulation of WTAP.

**Figure 2.**
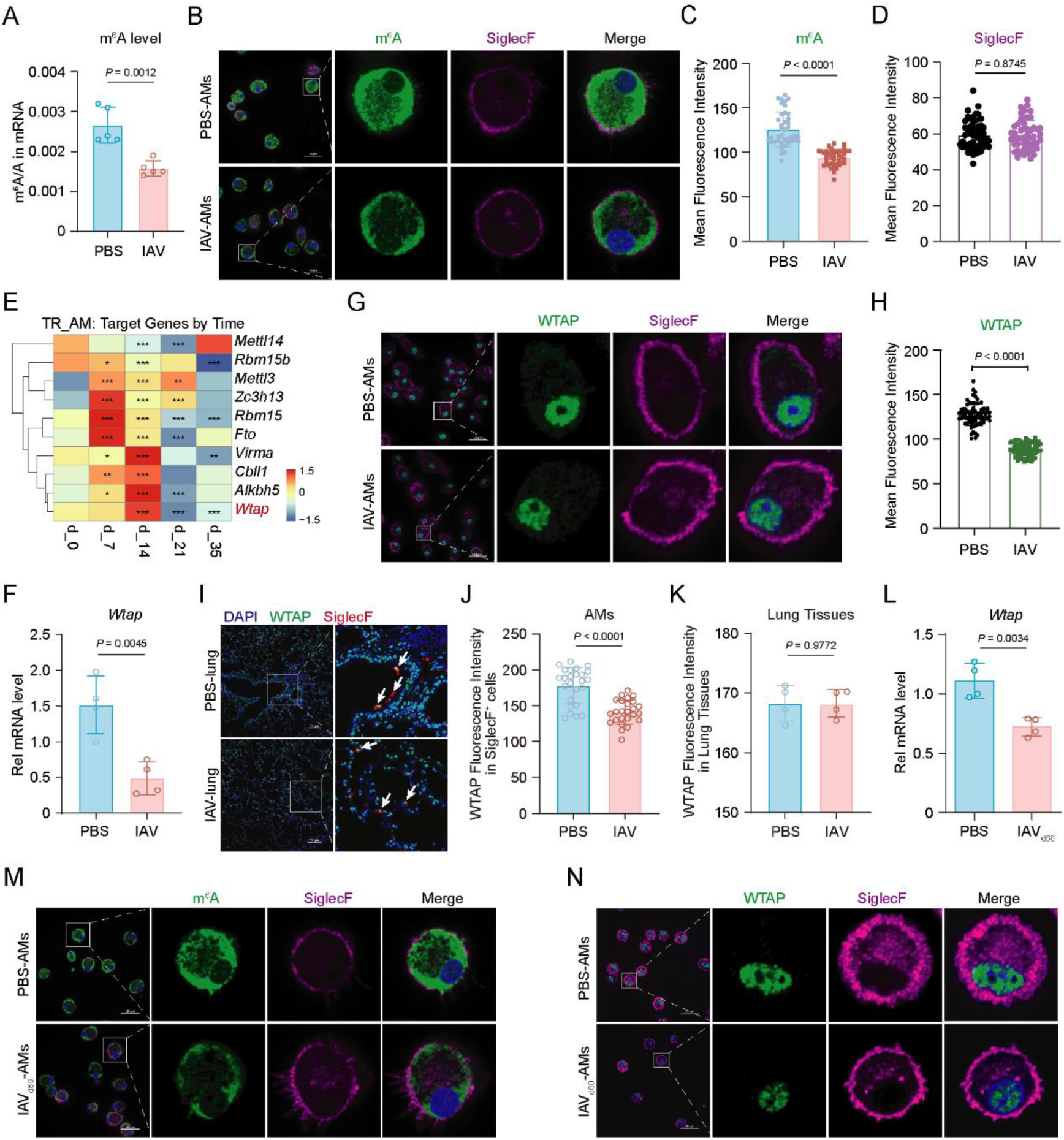
WTAP is downregulated in AMs during IAV-induced trained immunity. (**A**) LC–MS/MS quantification of m^6^A abundance in mRNA extracted from PBS- or IAV-AMs. (**B** to **D**) Representative imaging of m^6^A (green) and Siglec F (fuchsia) immunofluorescence staining in PBS-and IAV-AMs (B) and quantification of m^6^A (C) or Siglec F (D) intensity. (**E**) Heat maps showing the expression of m^6^A-related genes in TR_AMs from all time points. \**P* < 0.05, \*\**P* < 0.01, \*\*\**P* < 0.001. (**F**) qRT-PCR showing the mRNA abundance of *Wtap* in PBS- and IAV-AMs. n = 4 mice per group. (**G** and **H**) Representative imaging of WTAP (green) and Siglec F (fuchsia) immunofluorescence staining in PBS- and IAV-AMs (G) and quantification of WTAP intensity (H). Scale bar, 25 μm. (**I** to **K**) Representative confocal fluorescence imaging of lung slices from uninfected (PBS) and IAV-infected mice stained with antibodies against indicated proteins (I) and quantification of the mean fluorescence intensity of WTAP in Siglec F^+^ AMs (J) or in lung tissues (K). Scale bar, 100 μm. n = 4 mice per group. (**L**) qRT-PCR showing the mRNA abundance of *Wtap* in PBS- and IAV_d60_-AMs. n = 4 mice per group. (**M**) Representative imaging of m^6^A (green) and Siglec F (fuchsia) immunofluorescence staining in AMs from PBS- and day 60 IAV-infected (IAV_d60_) mice. (**N**) Representative imaging of WTAP (green) and Siglec F (fuchsia) immunofluorescence staining in AMs from PBS- and day 60 IAV-infected (IAV_d60_) mice. Data are presented as the mean ± s.d. in A, C, D, F, H and J-L with individual measurements overlaid as dots, and statistical analysis was performed in A, C, D, F, H and J-L using a two-tailed Student’s *t*-test or performed in E using Mann-Whitney U tests. Data in B, G, I, M and N are representative of three independent biological experiments.

### WTAP-deficient AMs exhibit enhanced phagocytic and metabolic capacity to protect mice against pulmonary bacterial infection

The above findings indicate that the reduced m^6^A abundance in accordance with down-regulated WTAP expression persists long-term in AMs from IAV-trained mice. This prompted us to investigate whether WTAP downregulation contributes to establishing a trained phenotype in these cells. To test this, we utilized *Wtap* conditional knockout mice (*Wtap*^fl/fl^ *LyzM*-Cre), previously generated by crossing *Wtap* flox mice (*Wtap*^fl/fl^) with myeloid lineage–specific *LyzM*-Cre mice (38). AMs purified from these mice showed nearly undetectable expression of WTAP (Supplementary Figure 6, A-C). We first assessed the phagocytic function of AMs isolated from *Wtap*^fl/fl^ and *Wtap*^fl/fl^ *LyzM*-Cre mice (referred to as WT and *Wtap*-cKO AMs hereafter) using multiple approaches (Figure 3A). RNA-seq analysis revealed that *Wtap*-cKO AMs significantly upregulated gene clusters related to phagocytosis, which are critical for macrophage antibacterial activity (Figure 3B and Supplementary Figure 6D). Moreover, peroxisome proliferator-activated receptor gamma (PPARγ), a well-known anti-inflammatory transcription factor (39), is predominantly expressed in AMs (40). It also plays a central role in regulating lipid metabolism and the expression of phagocytosis-associated receptors (41). We found genes involved in the PPARγ signaling pathway were also highly expressed in *Wtap*-cKO AMs (Supplementary Figure 6E). These data led us to hypothesize that WTAP deficiency in AMs enhances phagocytic capacity to ‘silently’ capture and clean microbial intruders.

**Figure 3.**
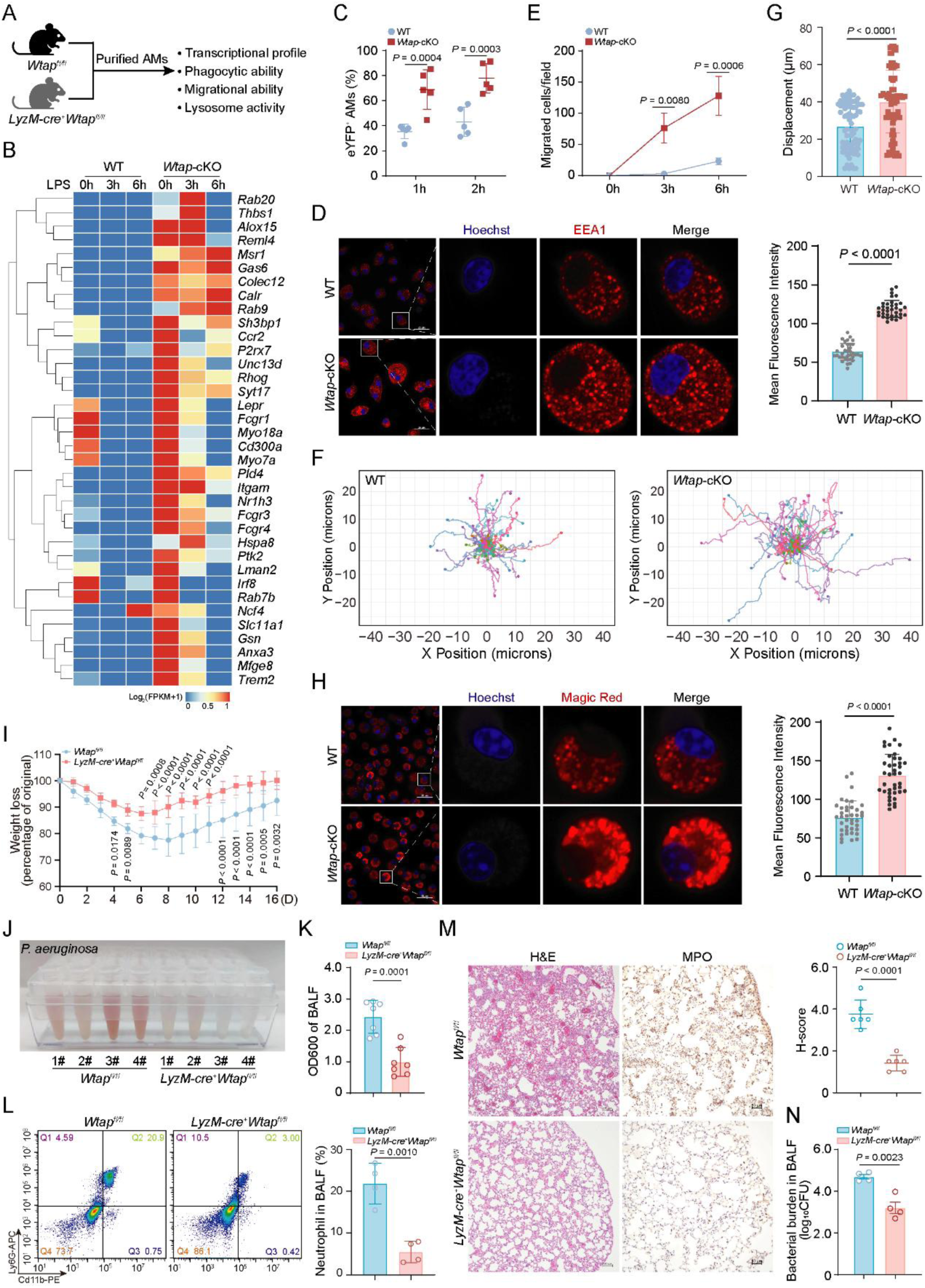
WTAP-deficient AMs exhibit enhanced phagocytic activity to protect mice against pulmonary bacterial infection. (**A**) Schema of the characteristic evaluation of WT and *Wtap*-cKO AMs purified respectively from *Wtap*^fl/fl^ and *LyzM*-Cre^+^ *Wtap*^fl/fl^ mice. (**B**) RNA-seq heat maps showing the mRNA abundance of genes related to phagocytosis. (**C**) Flow cytometry quantifying the phagocytosis of WT and *Wtap*-cKO AMs on eYFP-*E. coli*. (**D**) Representative imaging of EEA1 immunofluorescence staining in WT and *Wtap*-cKO AMs (left) and quantification of EEA1 intensity (right). Scale bar, 25 μm. (**E**) Quantification of the *in vitro* chemotaxis assay of WT and *Wtap*-cKO AMs. (**F**) Spider plot of randomly chosen WT and *Wtap*-cKO AMs, and colors represent individual AMs (n = 20 cells). (**G**) Quantification of net displacement (micrometers) of WT and *Wtap*-cKO AMs during a 1 h imaging session. (**H**) Representative imaging of Magic Red dye in WT and *Wtap*-cKO AMs (left) and quantification of Magic Red intensity (right). Scale bar, 25 μm. (**I**) Weight loss monitored over time of *Wtap*^fl/fl^ and *LyzM*-Cre^+^ *Wtap*^fl/fl^ mice infected with *P. aeruginosa*. n = 8 mice per group. (**J** and **K**) Observation of the bleeding (J) and measurement of the turbidity (K) in BALFs from *Wtap*^fl/fl^ and *LyzM*-Cre^+^ *Wtap*^fl/fl^ mice 12 hours after *P. aeruginosa* administration. n = 4 or 7 mice per group. (**L**) Flow cytometric plot shows Cd11b^+^ Ly6G^+^ neutrophils in BALFs from *Wtap*^fl/fl^ and *LyzM*-Cre^+^ *Wtap*^fl/fl^ mice at 12 h after *P. aeruginosa* infection (left) and quantification of the percentage of neutrophils (right). n = 4 mice per group. (**M**) Representative histology images of lung tissues from *Wtap*^fl/fl^ and *LyzM*-Cre^+^ *Wtap*^fl/fl^ mice at 12 h after *P. aeruginosa* infection (left) and H-score of H&E staining (right). Scale bar, 100 or 50 μm. N = 6 mice per group. (**N**) Quantification of bacterial burden of BALFs from *Wtap*^fl/fl^ and *LyzM*-Cre^+^ *Wtap*^fl/fl^ mice at 12 h after *P. aeruginosa* infection. n = 4 mice per group. Data are presented as the mean ± s.d. in C, D, E, G, H and K-N with individual measurements overlaid as dots, and statistical analysis was performed in C, E and I using Multiple Comparisons Following Two-Way ANOVA or performed in D, G, H and K-N using a two-tailed Student’s *t*-test. Data in D, H, J, L and M are representative of three independent biological experiments.

To prove this, we assessed the phagocytic capacity *Wtap*-cKO AMs. Incubation of AMs with eYFP-labeled *E. coli* followed by flow cytometric analysis revealed that *Wtap*-cKO AMs internalized a greater number of bacteria than WT AMs (Figure 3C). Consistent with this, immunofluorescence staining for the early endosome marker EEA1 demonstrated that more early phagosomes formed in *Wtap*-cKO AMs following *P. aeruginosa* infection (Figure 3D). Migration assays showed that although WT AMs could migrate toward *P. aeruginosa*, *Wtap*-cKO AMs exhibited a significantly enhanced migratory ability (Figure 3E and Supplementary Figure 6F). Accordingly, *Wtap*-cKO AMs displayed more pseudopodia (Supplementary Figure 6, G-H), and showed a significant increase in displacement compared to WT AMs (Figure 3, F and G, and Supplementary Figure 6I). These data collectively indicated that WTAP-deficient AMs possess a stronger ability to phagocytose bacteria.

Lysosomes serve as the central executors of phagocytosis in macrophages (42), containing abundant hydrolytic enzymes responsible for degrading macromolecules delivered via various membrane-trafficking pathways (43). We next assessed lysosomal activity in AMs. Co-incubation of WT and *Wtap*-cKO AMs with *P. aeruginosa* labeled with pHrodo red, a pH-sensitive dye that fluoresces only upon internalized vesicle acidification, revealed intensified red fluorescence in *Wtap*-cKO AMs, indicating increased lysosomal acidification upon WTAP deficiency (Supplementary Figure 6J). Meanwhile, Magic Red assay for cathepsin B activity showed enhanced lysosomal hydrolytic function in *Wtap*-cKO AMs (Figure 3H). Together, these data suggested that WTAP-deficient AMs exhibit enhanced phagocytic and hydrolytic capacity.

Given that AMs phagocytose inhaled bacteria to prevent alveolar inflammation (12), we hypothesized that enhancing their phagocytosis through WTAP downregulation should improve antibacterial defense. To test this, we employed a lung infection model of *P. aeruginosa*. Following a sublethal intratracheal challenge, *Wtap*^fl/fl^ *LyzM*-Cre mice exhibited significantly lower morbidity than *Wtap*^fl/fl^ controls (Figure 3I). Concurrently, severe BALF bleeding observed in WT controls was absent in *Wtap*^fl/fl^ *LyzM*-Cre mice (Figure 3, J and K, and Supplementary Figure 7A). Flow cytometry detections of BALF (Figure 3L) and the histopathological assessments of infected lungs (Figure 3M and Supplementary Figure 7B) further revealed reduced neutrophil infiltration, milder tissue damage, and less hemorrhage in *Wtap*^fl/fl^ *LyzM*-Cre mice. Consistent with enhanced host defense, bacterial burden in BALF was significantly lower in *Wtap*^fl/fl^ *LyzM*-Cre mice (Figure 3N and Supplementary Figure 7C). All these findings thus indicated that downregulation of WTAP enhances host defense against pulmonary bacterial infection by strengthening AM phagocytic and hydrolytic capacity.

WTAP deficiency drives an enhancement in phagocytic capacity within AMs, similar to prototypical TRIM induced by IAV (7). Cellular metabolism critically shapes the functional state of macrophages, including the induction, maintenance and regulation of TRIM (44). More importantly, the enhanced phagocytic and hydrolytic functions require a higher energy metabolism (45). So, we next explored whether the trained phenotype in WTAP-deficient AMs is accompanied by metabolic rewiring. As we expected, ATP levels measured with luciferin assays showed that deficiency of WTAP in AMs resulted in higher cellular ATP content (Figure 4A). RNA-Seq analysis further revealed that WTAP deficiency-induced TRIM is associated with upregulated expression of genes involved in the energetic metabolism, including glycolysis, tricarboxylic acid (TCA) cycle and lipid metabolism (Figure 4B and Supplementary Figure 8, A-C), consistent with enhanced ATP production. We then confirmed this metabolic shift through multiple approaches. TEM assays demonstrated a greater mitochondrial number in *Wtap*-cKO AMs relative to control cells (Figure 4, C and D). Since intracellular lipid droplets (LDs) serve as reservoirs of fatty acids for metabolic utilization, we next performed Oil Red O staining and observed increased LD accumulation in *Wtap*-cKO AMs (Figure 4, E and F), suggesting elevated lipid flux. In line with these findings, Seahorse metabolic flux analyses revealed significantly higher *ex vivo* oxygen consumption rates (OCRs) and extracellular acidification rates (ECARs) in *Wtap*-cKO AMs compared to WT controls (Figure 4, G-J), confirming a broad enhancement in metabolic activity.

**Figure 4.**
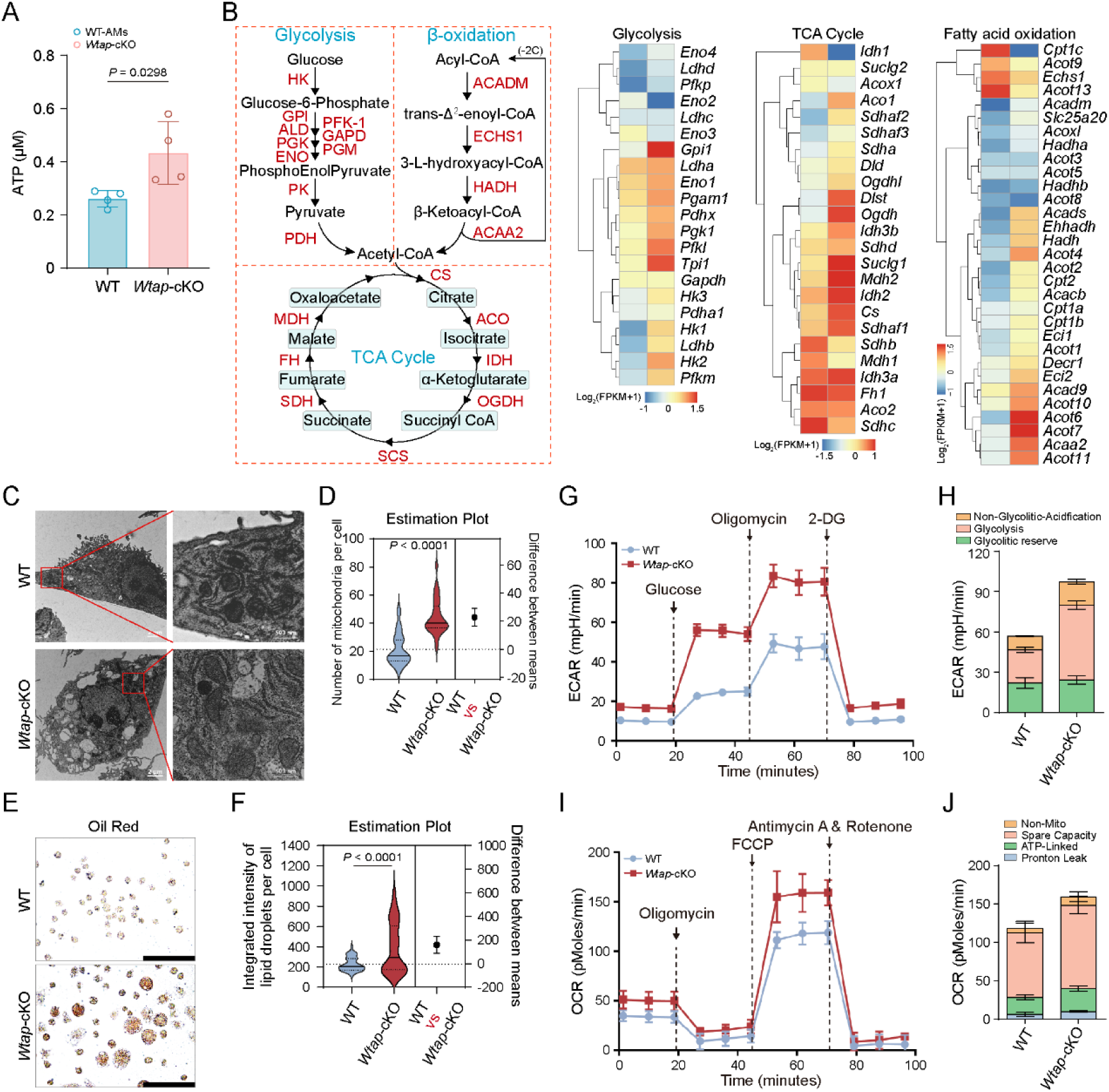
Metabolic rewiring in WTAP-deficient AMs enhance the characteristics of trained immunity. (**A**) Determination of ATP concentration in WT and *Wtap*-cKO AMs. (**B**) RNA-seq heat maps showing the mRNA abundance of enzyme genes involved in glycolysis, fatty acid oxidation and TCA cycle in WT and *Wtap*-cKO AMs. (**C** and **D**) Representative TEM micrograph of mitochondria in WT and *Wtap*-cKO AMs (C) and quantification of mitochondrion numbers (D). (**E** and **F**) Representative imaging of Oil Red staining in WT and *Wtap*-cKO AMs (E) and quantification of Oil Red intensity (F). (**G** and **H**) ECAR detection of WT and *Wtap*-cKO AMs in seahorse XF96 analyzer culture plates under basal conditions and then with sequential additions of various agonists and inhibitors at the indicated times (G), and quantification of baseline ECAR (H). (**I** and **J**) OCR detection of WT and *Wtap*-cKO AMs in seahorse XF96 analyzer culture plates under basal conditions and then with sequential additions of various agonists and inhibitors at the indicated times (I), and quantification of baseline OCR (J). Data are presented as the mean ± s.d. in A with individual measurements overlaid as dots, and statistical analysis was performed in A, D and F using a two-tailed Student’s *t*-test. Data in C and E are representative of three independent biological experiments.

Together, these results indicated that WTAP-deficient AMs exhibit enhanced phagocytic capacity fueled by enhanced metabolic adaptation to effectively protect mice against pulmonary bacterial infection.

### WTAP regulates AM trained immunity through m^6^A modification

Since WTAP participates in establishing IAV-induced TRIM in AMs, we next sought to elucidate the underlying regulatory mechanism. As a key regulatory subunit of the m^6^A methyltransferase complex, WTAP is essential for anchoring and stabilizing METTL3 and METTL14 to nuclear speckles, thereby regulating m^6^A modification of RNA (46). To determine whether WTAP affects AM TRIM via m^6^A, we first compared global m^6^A levels in mRNAs from WT and *Wtap*-cKO AMs using LC-MS/MS. The results showed a marked reduction in *Wtap*-cKO AMs (Figure 5A), a trend further supported by immunofluorescence analysis (Supplementary Figure 9, A and B). We then performed direct RNA sequencing using the Oxford Nanopore Technologies platform (ONT DRS) on mRNAs from WT and *Wtap*-cKO AMs. By detecting modified adenosines in individual DRS reads, we identified 107,576 and 15,553 high-confidence m^6^A sites in the WT and *Wtap*-cKO AMs, respectively (Figure 5B). These sites predominantly contained the canonical ‘RRACH (R=G or A; H=A, C or U)’ motif (Supplementary Figure 9C) and were enriched in CDS and 3’ UTR regions, as expected for m^6^A (Supplementary Figure 9D). Notedly, WTAP deficiency significantly reduced the modification abundance at the 3’ UTR region near the stop codon (Figure 5C). Approximately 11,631 m^6^A sites across 2,900 genes showed significantly decreased methylation in *Wtap*-cKO AMs (Figure 5D). Pathway enrichment analysis revealed that these identified genes were associated with phagocytosis, bacterial infection, innate immune responses, and glucose metabolism (Figure 5E and Supplementary Figure 9E). Moreover, more genes were upregulated in *Wtap*-cKO AMs (Supplementary Figure 9F), particularly those involved in phagocytosis and innate immune responses (fig. S9G). We further identified 318 overlapping transcripts that exhibited both reduced m^6^A marks and increased mRNA abundance after WTAP deletion (Supplementary Figure 9H). KEGG enrichment analyses confirmed their strong association with phagocytosis, infection, immunity, and glucose metabolism (Supplementary Figure 9I). These results indicated that WTAP shapes the TRIM phenotype in AMs by regulating the m^6^A modification of mRNAs related to phagocytotic and metabolic functions.

**Figure 5.**
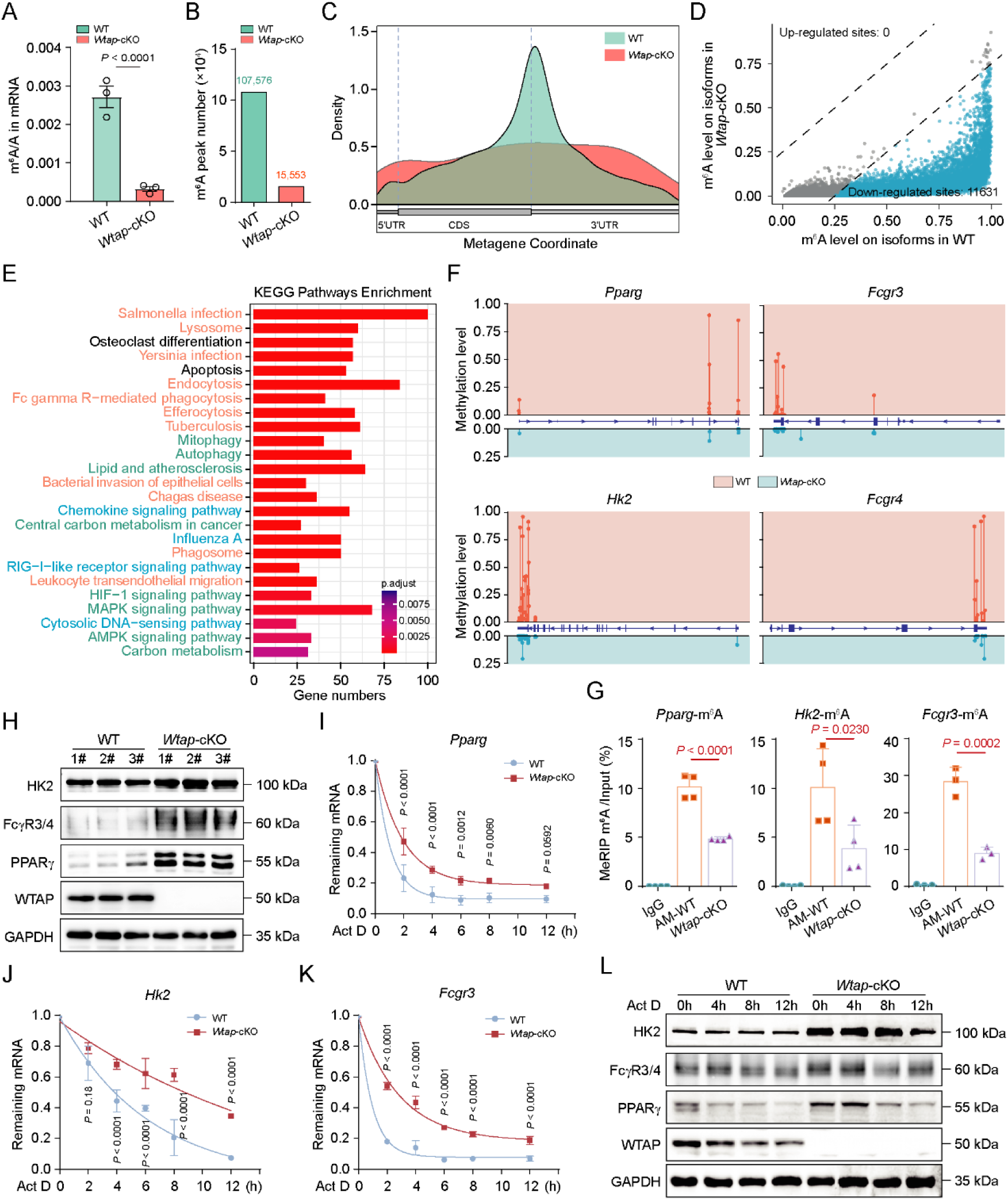
WTAP regulates AM trained immunity through m^6^A modification. (**A**) LC–MS/MS quantification of m^6^A abundance in mRNA extracted from WT and *Wtap*-cKO AMs. (**B**) Statistics of m^6^A modification sites in WT and *Wtap*-cKO AMs. (**C**) Metagene profiles of m^6^A peak distribution across the 5’ UTR, CDS, and 3’ UTR in WT and *Wtap*-cKO AMs. (**D**) Scatter plot showing the differentially m^6^A sites in *Wtap*-cKO AMs compared with WT controls. (**E**) KEGG enrichment analysis of the genes with decreased m^6^A marks in *Wtap*-cKO AMs compared with WT controls. (**F**) Genome browser screenshot showing WTAP-dependent m^6^A sites in the *Pparg*, *Hk2*, *Fcgr3*, and *Fcgr4* transcripts. (**G**) MeRIP-qPCR showing the abundance of *Pparg*, *Hk2* and *Fcgr3* transcripts in WT and *Wtap*-cKO AMs. (**H**) Immunoblotting analysis of the expression of PPARγ, HK2 and FcγR3 in WT and *Wtap*-cKO AMs. (**I** to **K**) qRT–PCR showing the mRNA abundance of *Pparg*, *Hk2* and *Fcgr3* in WT and *Wtap*-cKO AMs treated with Act D at the indicated time points. (**L**) Immunoblotting analysis of the expression of PPARγ, HK2 and FcγR3 in WT and *Wtap*-cKO AMs treated with Act D at the indicated time points. Data are presented as the mean ± s.d. in A, G and I-K with individual measurements overlaid as dots, and statistical analysis was performed in performed in A and G using a two-tailed Student’s *t*-test or performed in I-K using Multiple Comparisons Following Two-Way ANOVA. Data in H and L are representative of three independent biological experiments.

To verify this mechanism, we then evaluated the identified WTAP target genes involved in the phagocytotic and metabolic functions of AMs, including PPARγ, HK2, FcγR3 and FcγR4. PPARγ is a nuclear transcription factor, usually binding to PPAR-responsive regulatory elements in the genome to modulate the expression of genes involved in fatty acid oxidation, inflammation and phagocytosis in AMs (47). Hexokinase 2 (HK2) is a rate-limiting glycolytic enzyme that controls glycolytic flux and mitochondrial activity (48). Fcγ receptors (FcγR) mediate opsonic phagocytosis of IgG-opsonized pathogens (49). All these genes play a significant role in the maintenance and remodeling of the lung homeostasis function of AMs. ONT DRS data revealed high-confidence and WTAP-sensitive m^6^A sites in the 3’ UTR near the stop codon of *Hk2*, *Pparg*, *Fcgr3* and *Fcgr4* transcripts (Figure 5F). We then designed a gene-specific primer pair to measure the change in m^6^A abundance of these transcripts via MeRIP-qPCR assay. The results confirmed a significant decrease in m^6^A marks on *Hk2*, *Pparg*, and *Fcgr3* mRNAs in *Wtap*-cKO AMs compared with control cells (Figure 5G). Furthermore, as the mRNA methylation level decreased, the abundance of the PPARγ, HK2 and FcγR3 protein in *Wtap*-cKO AMs significantly increased (Figure 5H). Hence, the detected m^6^A marks in *Hk2*, *Pparg* and *Fcgr3* transcripts are direct substrates of WTAP and are crucial for regulating their protein output.

Given that m^6^A predominantly promotes mRNA decay (50, 51), we then assessed the mRNA stability of the *Hk2*, *Pparg* and *Fcgr3* transcripts through RNA decay assays. The results showed that the decay rate of these transcripts was considerably slower in *Wtap*-cKO AMs when transcription was halted with actinomycin D (Act D) (Figure 5, I and K), and the turnover of their corresponding proteins followed the same trend (Figure 5L). Thus, WTAP directly targets *Hk2*, *Pparg* and *Fcgr3* mRNAs for m^6^A modification to promote their degradation. Accordingly, reanalyzing the aforementioned scRNA-seq data [accession nos. GSE208294] revealed that the trained cluster 0 TR_AMs also exhibited downregulated *Wtap* during recovery from IAV infection (Supplementary Figure 10A). This decrease was accompanied by significant upregulation of phagocytic genes, including the direct targets (*Hk2*, *Pparg*, *Fcgr3*, *Fcgr4*), as well as other related genes (Supplementary Figure 10B). Correlation analysis further supported a strong negative relationship between *Wtap* and its target genes (Supplementary Figure 10, C and D). Collectively, these findings underscored the role of WTAP-dependent m^6^A methylation in regulating the phagocytotic and metabolic function, thereby affecting the establishment of TRIM in AMs.

### Trained AMs associated with lower expressed WTAP alleviate the severity of pulmonary diseases

Having established the antibacterial function of IAV-trained, low-WTAP AMs in mice, we then investigated whether a counterpart population exists in human BALF and is linked to clinical disease severity. The analysis was performed using published data on scRNA-seq in BALF from four healthy individuals (HC1–HC4) and nine patients with mild (M1–M3) and severe (S1–S6) COVID-19 infection [accession nos. GSE145926] (52). Clustering analysis showed 7 major cell types composed of macrophages, neutrophils, dendritic cells (DCs), natural killer (NK) cells, T cells, B cells and epithelial cells, identified by signature genes (Figure 6A). Among these, the macrophage subpopulation was the predominant subset (Figure 6A). We next analyzed the *WTAP* expression in BAL macrophages from healthy individuals, mild and severe COVID-19 patients and found that mild patients have lower expression of *WTAP* compared to healthy individuals and severe patients (Figure 6, B and C). This expression trend of *WTAP* was confirmed in all the samples (Figure 6D). In contrast, the expression of *HK2*, *PPARG* and *FCGR3A* was correspondingly and significantly upregulated (Figure 6E), indicating a negative correlation with WTAP expression. We next assessed how WTAP-mediated functional remodeling of BAL macrophages shapes COVID-19 severity. KEGG enrichment analysis showed that, compared to healthy individuals, the upregulated genes in BAL macrophages from mild COVID-19 patients enriched in the pathways related to phagocytosis, chemokine signaling, antigen processing and presentation (Figure 6F), which is consistent with the characteristics of the low WTAP-mediated trained AMs. Furthermore, compared to severe cases, BAL macrophages in patients with mild COVID-19 displayed a trained immunity phenotype characterized by enhanced capacities for phagocytosis, antigen presentation, and glucose metabolism (Figure 6G). Hence, the establishment of low WTAP-mediated TRIM establishment in AMs is a key determinant of disease severity in respiratory viral infections, including influenza and COVID-19.

**Figure 6.**
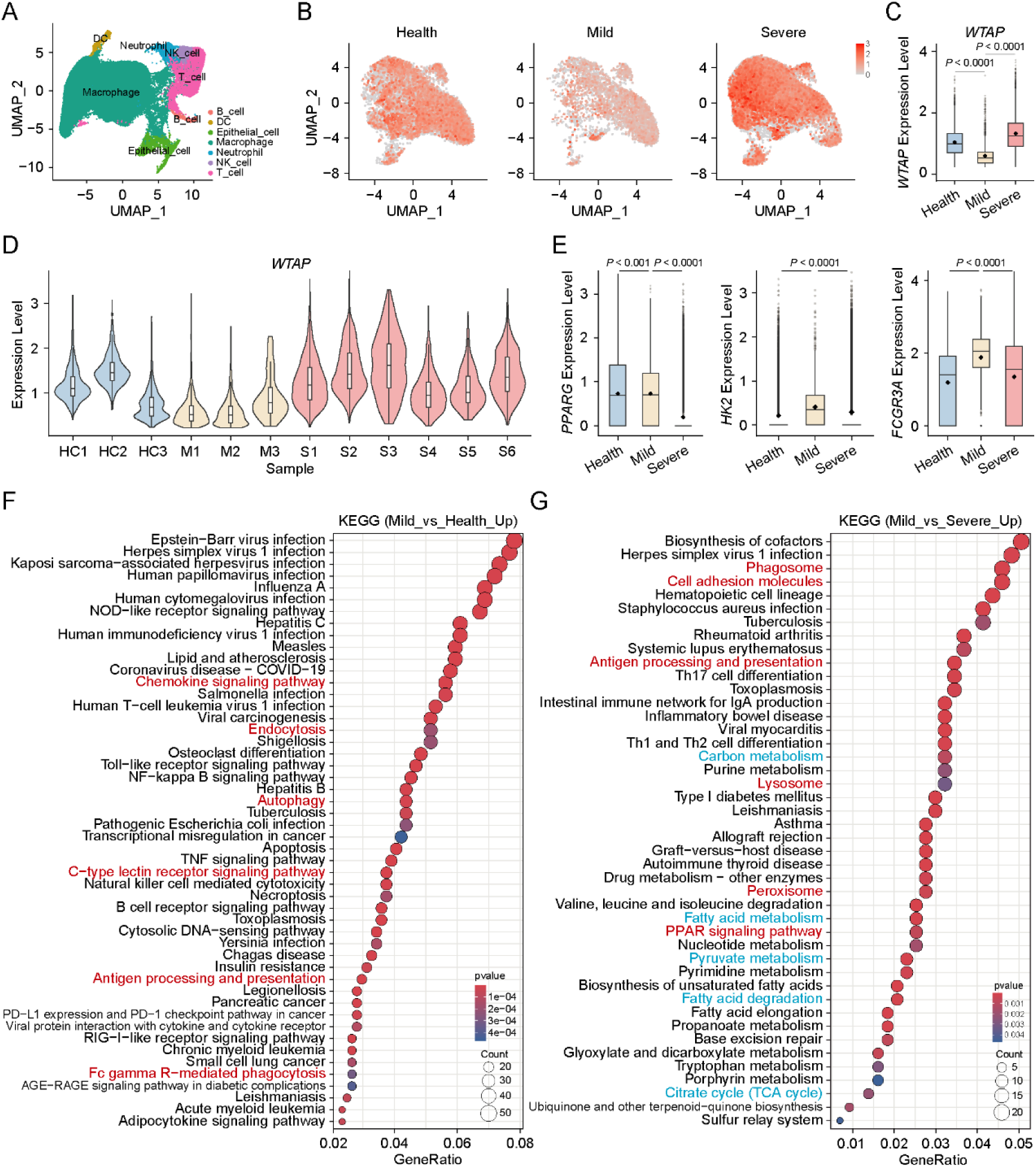
Trained AMs associated with lower expressed WTAP alleviate the severity of pulmonary diseases. (**A**) UMAP presentation of major cell types and associated clusters in BALFs (n = 13, including three patients with mild COVID-19 (M1–M3), six patients with severe/critical infection (S1–S6) and four healthy controls (HC1–HC4)). (**B**) UMAP projection of the expression of *WTAP* in BAL macrophages from healthy controls (HC) and patients with mild (M) or severe (S) COVID-19 infection. (**C**) Box plot showing the mRNA expression of *WTAP* in BAL macrophages from healthy controls (HC) and patients with mild (M) and severe (S) COVID-19 infection. (**D**) Expression of *WTAP* by BALF macrophages from each sample. (**E**) Box plot showing the mRNA expression of *PPARG*, *HK2* and *FCGR3A* in BAL macrophages from healthy controls (HC) and patients with mild (M) and severe (S) COVID-19 infection. (**F** and **G**) KEGG enrichment analysis of the upregulated genes in BAL macrophages from mild COVID-19 patients versus healthy controls (F) or severe cases (G). Statistical analysis was performed in C and E using Mann-Whitney U test.

The phagocytic function of AMs is a central mechanism not only in host defense but also in the development of chronic pulmonary diseases such as asthma, chronic obstructive pulmonary disorder (COPD) and pulmonary alveolar proteinosis (PAP) (53). COPD is a major public health problem and AMs from patients with COPD are defective in phagocytosing bacteria or apoptotic epithelial cells (54). We next investigated the potential role of WTAP in regulating AM phagocytosis during COPD pathogenesis. The scRNA-seq dataset [accession nos. GSE136831] (55) in distal lung parenchyma samples obtained from healthy donors and patients with COPD were analyzed, and identifying 39 discrete cell types based on distinct markers with no discernible batch (Supplementary Figure 11A). Cluster analysis of AMs revealed 7 distinct subpopulations (Supplementary Figure 11B). Notably, *WTAP* expression was significantly elevated in nearly all AM clusters from COPD patients compared to healthy controls (Supplementary Figure 11, C and D). Moreover, following WTAP expression rose in AMs from COPD patients (Supplementary Figure 11E), the expression of its target genes (*PPARG*, *FCGR3A* and *FCGR3B*) became significantly lower than those in healthy individuals (Supplementary Figure 11F). Further KEGG enrichment of the downregulated genes in AMs from COPD patients compared to healthy controls returned biological features including efferocytosis, lysosomal hydrolysis, and several carbon and fatty acid metabolic pathways (Supplementary Figure 11G), consistent with WTAP-mediated functional characteristics. Hence, upregulation of WTAP in AMs drives phagocytic defects in COPD patients by suppressing target genes (e.g., PPARG, FCGR3A and FCGR3B) and downregulating associated functional pathways.

Together, all these data indicated that high expression of WTAP in the disease state drives phagocytic impairment in AMs, which in turn contributes to aggravated disease severity in both severe COVID-19 and COPD.

### Inhibition of m^6^A modification enhances antibacterial activity

RNA methyltransferases are important regulators of RNA sensing and innate immune activation and represent novel immune-regulatory targets (56–58). For example, as an oral small molecule, STC-15 is the first RNA methyltransferase inhibitor to enter clinical development (59). Inhibition of METTL3 by STC-15 leads to prominent upregulation of genes associated with innate immunity, and may establish durable antitumor immune memory in mice (30). We have found that low m^6^A modification caused by downregulated WTAP can enhance the phagocytosis of AMs, so we further verified whether the METTL3 inhibitor STC-15 can also enhance this characteristic of AMs by reducing the m^6^A modification. The results showed that reducing the abundance of m^6^A marks with STC-15 (Figure 7A) could observably enhance the phagocytosis of AMs to bacteria (Figure 7B and Supplementary Figure 12A), and more early phagosomes were formed in STC-15-pretreated AMs compared with control cells (Figure 7, C and D). With the enhancement of phagocytic ability, the enzymatic activity of cathepsin B in lysosomes from STC-15-pretreated AMs also significantly increased (Figure 7, E and F). In addition, chemotaxis assays showed STC-15-pretreated AMs had a marked promotion in their ability to migrate toward *P. aeruginosa* (Figure 7G and Supplementary Figure 12B). These functional enhancements were supported by metabolic flux, as evidenced by a significant increase in intracellular ATP content in STC-15-pretreated cells (Figure 7H).

**Figure 7.**
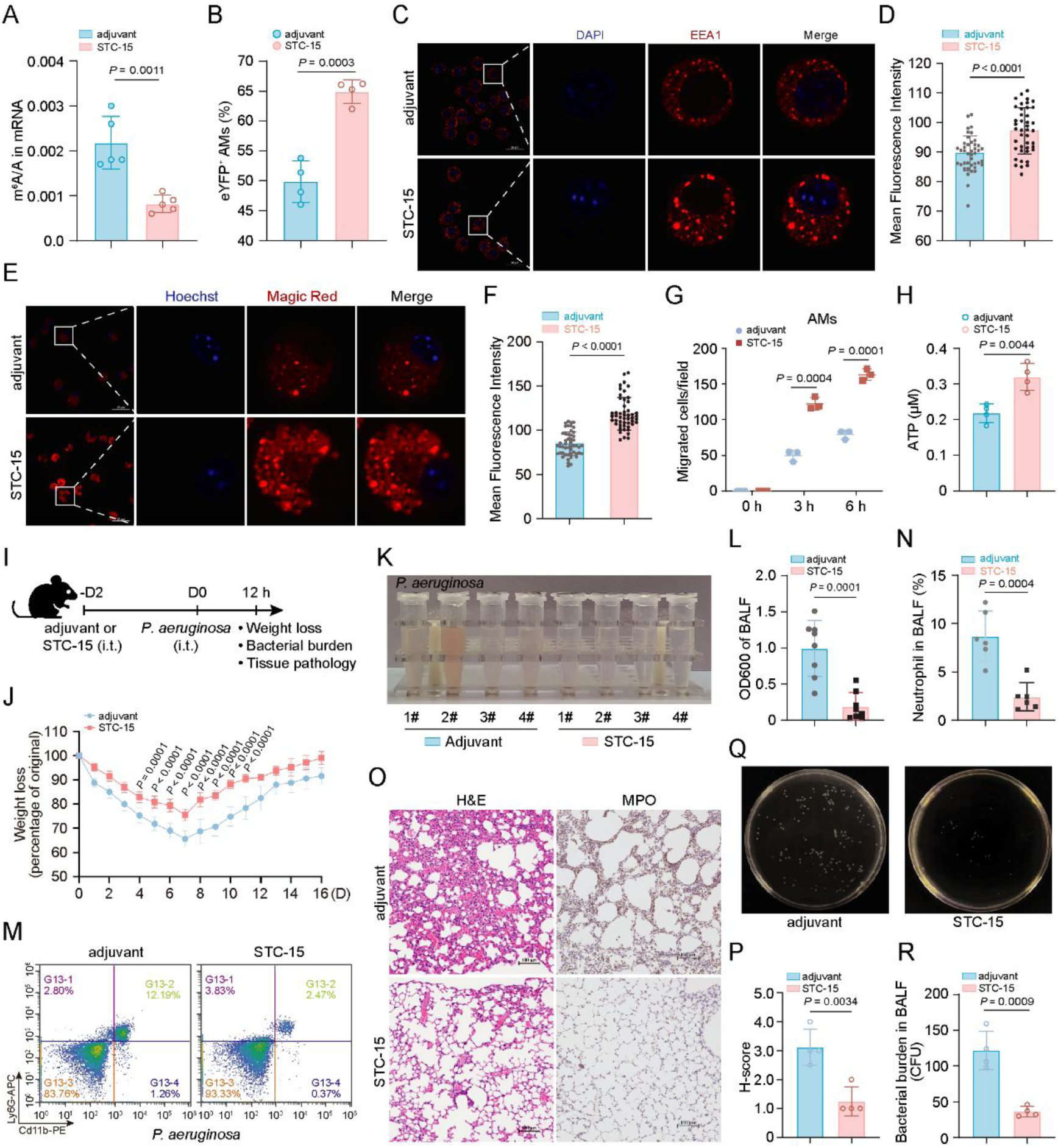
Inhibition of m^6^A modification enhances antibacterial activity. (**A**) LC–MS/MS quantification of m^6^A abundance in mRNA extracted from AMs treated with adjuvant or STC-15. (**B**) Flow cytometry quantifying the phagocytosis of adjuvant- or STC-15-pretreated AMs on eYFP-*E. coli*. (**C** and **D**) Representative imaging of EEA1 immunofluorescence staining in adjuvant- or STC-15-pretreated AMs (C) and quantification of EEA1 intensity (D). Scale bar, 25 μm. (**E** and **F**) Representative imaging of Magic Red dye in adjuvant- or STC-15-pretreated AMs (E) and quantification of Magic Red intensity (F). Scale bar, 25 μm. (**G**) Quantification of the *in vitro* chemotaxis assay of adjuvant- or STC-15-pretreated AMs. (**H**) Determination of ATP concentration in AMs from adjuvant- and STC-15-pretreated mice. n = 4 mice per group. (**I**) Experimental schema of i.t. STC-15 pretreatment in mice followed by i.t. *P. aeruginosa* infection. (**J**) Weight loss monitored over time of adjuvant- and STC-15-pretreated mice infected with *P. aeruginosa*. n = 8 mice per group. (**K** and **L**) Observation of the bleeding in (K) and measurement of the turbidity of (L) BALFs from adjuvant- and STC-15-pretreated mice 12 hours after *P. aeruginosa* administration. n = 4 or 8 mice per group. (**M** and **N**) Flow cytometric plot shows Cd11b^+^ Ly6G^+^ neutrophils in BALFs from adjuvant- or STC-15-pretreated mice at 12 h after *P. aeruginosa* infection (M) and quantification of the percentage of neutrophils (N). n = 6 mice per group. (**O** and **P**) Representative histology images of lung tissues from adjuvant- or STC-15-pretreated mice at 12 h after *P. aeruginosa* infection (O) and H-score of H&E staining (P). Scale bar, 100 μm. n = 4 mice per group. (**Q** and **R**) Representative bacterial plate pictures of BALFs from adjuvant- or STC-15-pretreated mice at 12 h after *P. aeruginosa* infection (Q) and quantification of bacterial burden (R). n = 4 mice per group. Data are presented as the mean ± s.d. in A, B, D, F-H, L, N, P and R with individual measurements overlaid as dots, and statistical analysis was performed in performed in A, B, D, F, H, L, N, P and R using a two-tailed Student’s *t*-test or performed in G and J using Multiple Comparisons Following Two-Way ANOVA. Data in C, E, K, M, O and Q are representative of three independent biological experiments.

In order to confirm that these changes are caused by the fluctuant m^6^A modification of the *Hk2*, *Pparg* and *Fcgr3/4* transcripts, we measured the change in m^6^A abundance of these transcripts via MeRIP-qPCR assay. The results revealed a marked decrease in m^6^A enrichment on these transcripts in STC-15-treated AMs compared to controls (Supplementary Figure 12C). With the decrease of the m^6^A abundance, the mRNA stability of these transcripts showed a significant increase (Supplementary Figure 12, D-F), as well as the abundance of the PPARγ, HK2 and FcγR3/4 protein (Supplementary Figure 12, G and H). All these data indicated that reduced m^6^A modification caused by both genetic (*Wtap*-cKO) and pharmacological (METTL3 inhibitor) treatment underlies the enhanced phagocytic and hydrolytic activity in AMs. This functional boost is achieved through the stabilization of key metabolic and phagocytic transcripts, including *Hk2*, *Pparg*, and *Fcgr3/4*.

To further verify the effect of reducing m^6^A modification of AMs on resistance to pulmonary bacterial infection in mice, we administered the vehicle or STC-15 (10 mg/Kg) intratracheally to wild-type (WT) C57BL/6 mice, and at 2 d after instillation, mice were infected intratracheally with *P. aeruginosa* (Figure 7I). Upon intratracheal infection of *P. aeruginosa* with a sublethal dose for 12 h, STC-15-primed mice lost less body weight compared to controls (Figure 7J). Similarly, severe bleeding in BALF was not observed in STC-15-primed mice compared with control mice (Figure 7K and Supplementary Figure 12I), which was corroborated with decreased exuded proteins and infiltrated neutrophils in BALF (Figure 7, L-N). Pathologically, H&E staining revealed a significant decrease in inflammatory cell infiltration and lung tissue edema in STC-15-primed group, which was consistent with the lung injury score results (Figure 7, O and P, and Supplementary Figure 12J). Accordingly, STC-15-primed mice had lower bacterial burden in their BALF from lungs than did control mice (Figure 7, Q and R). Thus, STC-15-primed mice were significantly less susceptible to *P. aeruginosa* with controlled pulmonary infection better than control mice.

### LNP-delivered *Wtap*-siRNA delivery can protect mice from bacterial pneumonia

To target the WTAP/m^6^A axis in AMs more effectively and precisely, we used lipid nanoparticle (LNP) delivery technology. LNPs represent an efficient and biocompatible platform for genetic drug delivery by overcoming various extracellular and intracellular barriers (60). To further enhance specificity, LNPs can be functionalized with targeting ligands that bind cognate receptors on desired cell types, promoting receptor-mediated endocytosis (61). For example, mannose-conjugated PEG lipids on LNPs facilitate RNA delivery to CD206-high macrophages (e.g., Kupffer cells and AMs) through specific interaction with the mannose receptor (62). To assess the therapeutic potential of this approach in bacterial pneumonia, we formulated mannose-decorated LNPs loaded with *Wtap*-siRNA and administered the resulting si*Wtap*-LNPs (0.5 mg/Kg) intratracheally to C57BL/6 mice (Figure 8A). At 2 days post-instillation, mice were euthanized and AMs were isolated for WTAP expression analysis. The results showed that WTAP expression was significantly suppressed in AMs from si*Wtap*-LNP-pretreated mice (Figure 8, B and C), whereas its expression in lung parenchyma tissue remained unaffected (Figure 8D). Furthermore, a significant negative correlation was observed between WTAP and its target genes (Figure 8E). These data indicated that mannose-decorated LNPs can selectively and effectively deliver siRNA to AMs. So, we next adopted a lung infection model of *P. aeruginosa* in LNP-pretreated mice. Upon intratracheal infection of *P. aeruginosa* with a sublethal dose for 12 h, severe bleeding in BALF was not observed in si*Wtap*-LNP-primed mice compared with control mice (Figure 8, F and G), which was corroborated with decreased exuded proteins (Figure 8H). Pathologically, flow cytometry and H&E staining revealed a significant decrease in inflammatory cell infiltration and lung tissue edema in si*Wtap*-LNP-primed group (Figure 8, I and J). Accordingly, mice primed with si*Wtap*-LNP showed a reduced bacterial burden in lung BALF compared to control mice (Figure 8K). All these data thus suggested that LNP-mediated siRNA delivery targeting AM WTAP represents a potential novel strategy for the prevention and treatment of bacterial pneumonia.

**Figure 8.**
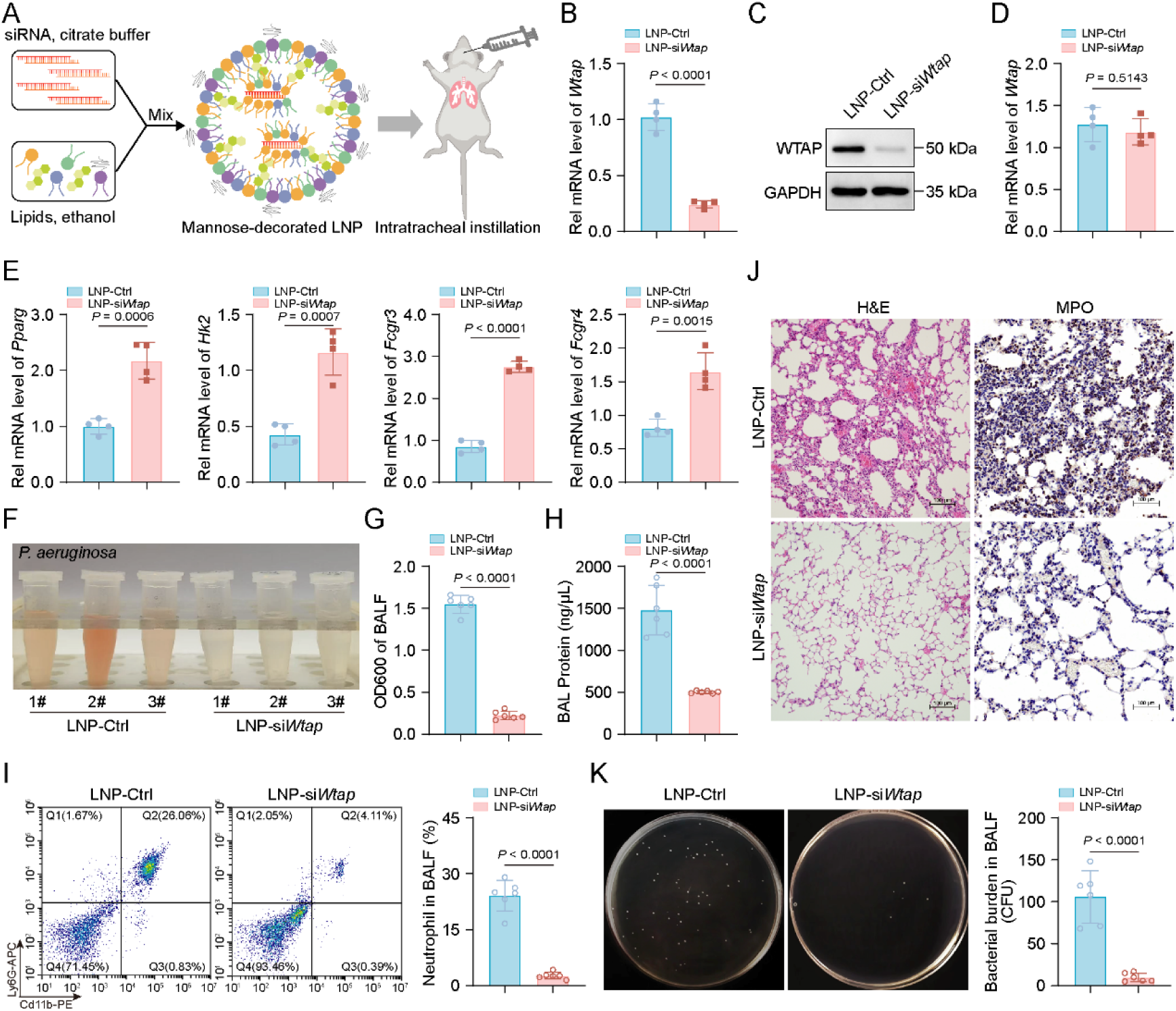
LNP-delivered Wtap-siRNA delivery can protect mice from bacterial pneumonia. (**A**) Schematic illustration of Mannose-decorated LNP synthesis and instillation. (**B** and **C**) qRT-PCR (B) and Immunoblotting (C) analysis of WTAP expression in AMs from LNP-Ctrl- or LNP-*siWtap*-pretreated mice. n = 4 mice per group. (**D**) qRT-PCR analysis of WTAP expression in lung tissues from LNP-Ctrl- or LNP-*siWtap*-pretreated mice. n = 4 mice per group. (**E**) qRT-PCR analysis of *Pparg*, *Hk2*, *Fcgr3* and *Fcgr4* in AMs from LNP-Ctrl- or LNP-*siWtap*-pretreated mice. n = 4 mice per group. (**F** and **G**) Observation of the bleeding in (F) and measurement of the turbidity of (G) BALFs from LNP-Ctrl- or LNP-*siWtap*-pretreated mice 12 hours after *P. aeruginosa* administration. n = 3 or 6 mice per group. (**H**) Quantification of BAL protein from LNP-Ctrl- or LNP-*siWtap*-pretreated mice 12 hours after *P. aeruginosa* administration. n = 6 mice per group. (**I**) Flow cytometric plot shows Cd11b^+^ Ly6G^+^ neutrophils in BALFs from LNP-Ctrl- or LNP-*siWtap*-pretreated mice at 12 h after *P. aeruginosa* infection (left) and quantification of the percentage of neutrophils (right). n = 6 mice per group. (**J**) Representative histology images of lung tissues from LNP-Ctrl- or LNP-*siWtap*-pretreated mice at 12 h after *P. aeruginosa* infection. (**K**) Representative bacterial plate pictures of BALFs from LNP-Ctrl- or LNP-*siWtap*-pretreated mice at 12 h after *P. aeruginosa* infection (left) and quantification of bacterial burden (right). n = 6 mice per group. Data are presented as the mean ± s.d. in B, D, E, G-I and K with individual measurements overlaid as dots, and statistical analysis was performed using a two-tailed Student’s *t*-test. Data in C, F and I-K are representative of three independent biological experiments.

## Discussion

Secondary bacterial pneumonia following influenza remains a major clinical challenge, largely attributable to the depletion and functional impairment of AMs. Consequently, therapeutic strategies aimed at functionally reprogramming AMs hold significant promise for preventing life-threatening post-viral complications. In this study, we elucidate a previously unknown epitranscriptomic mechanism governing the establishment of TRIM in AMs. IAV-training persistently downregulates expression of the m^6^A ‘writer’ protein WTAP in AMs, leading to reduced m^6^A methylation on key phagocytic and metabolic transcripts such as *Pparg*, *Hk2*, *Fcgr3* and *Fcgr4*. This epitranscriptomic alteration enhances mRNA stability and subsequent protein expression of these genes, thereby consolidating TRIM phenotype and boosting bacterial clearance. Ultimately, this RNA-centered reprogramming provides prolonged defense against secondary bacterial infections, representing a novel mechanism of innate immune memory in tissue-resident macrophages.

Traditionally, TRIM has been characterized by epigenetic and metabolic rewiring. Our findings position epitranscriptomic modification, specifically m^6^A methylation, as another fundamental pillar governing this process. This expands the mechanistic understanding of how short-lived RNA modifications can mediate long-term functional adaptation. Specifically, by modulating the expression of key epitranscriptomic regulators, RNA modification in nascent mRNAs can be continuously adjusted and harnessed to induce long-lasting phenotypic remodeling, which is a dimension that has remained underexplored. Furthermore, our work reveals the dynamic complexity of the mechanism for TRIM. Previous study has indicated that influenza-induced monocyte-derived AMs confer prolonged antibacterial protection primarily through increased interleukin-6 production (6). In contrast, our study reveals that resident AMs sustain long-term antibacterial defense by enhancing phagocytosis and metabolic capability through epitranscriptomic modification mediated by WTAP-m^6^A axis. Thus, rather than activating a universal trained state, the same stimulus can imprint distinct memories that generate specialized functional phenotypes in resident versus recruited macrophage populations.

Phagocytosis represents an essential functional pillar of the innate immunity (42, 63), and constitutes the lung’s preferred initial response to invasive pathogens and inhaled particulates (64). AMs patrol the alveolar spaces and phagocytose bacterial invaders, thereby helping to prevent damaging inflammation (12). A key mechanism that increases susceptibility to secondary bacterial infection during influenza is the suppression of AM phagocytosis by lung T cell-derived IFN-γ (65). Given the critical role of AM phagocytosis in resolving lung injury and preventing secondary infection, reshaping this function represents a promising strategy to counter viral-bacterial synergy and resolve tissue inflammation. Our work reveals that a prolonged TRIM driven by m^6^A hypomethylation in AMs can counteract this vulnerability and protect against secondary infection. This WTAP/m^6^A axis programs a unique form of TRIM that specifically enhances phagocytic capacity without increasing inflammatory output. By selectively targeting WTAP in AMs using mannose-functionalized LNP-delivered siRNA, we achieved significant protection against secondary bacterial pneumonia in a murine model. Thus, targeting WTAP in AMs offers a precise and translatable therapeutic strategy for clinical development. Most significantly, by enhancing macrophage function instead of targeting pathogens directly, this host-directed strategy not only bypasses antibiotic resistance but also offers potential broad-spectrum protection against diverse respiratory infections.

The dysregulation of WTAP expression was also observed in various lung diseases, such as patients with severe COVID-19, COPD or acute lung injury (38), and these changes are significantly correlated with lung homeostasis and disease prognosis. Importantly, impaired phagocytosis of AMs and accompanying bacterial infections is a typical characteristic of many chronic lung diseases like COPD (66), PAP (67) and asthma (68). Our data analyses confirm a significant correlation between elevated WTAP expression and impaired phagocytic activity in AMs from patients with severe COVID-19 or COPD. Thus, the WTAP/m^6^A axis serves not only as a therapeutic target but also as a source of biomarkers in these lung diseases; dynamic changes in m^6^A modification and WTAP expression can monitor disease progression and inform prognosis.

TRIM holds significant therapeutic potential in immunosuppressed populations, including those with cancer or infections, yet it also serves as a key driver of chronic inflammatory and autoimmune disorders (69, 70). For example, a TRIM signature in systemic lupus erythematosus (SLE) patients characterized by HSC-skewed myeloid lineage reprogramming may exacerbate immune responses and flares (71). Similarly, systemic inflammation triggered by periodontitis can epigenetically reprogram HSCs to drive myeloid overproduction and worsen inflammatory arthritis (72). Autoimmune manifestations following SARS-CoV-2 infection and vaccination further underscore this link (73). Thus, a major hurdle in harnessing TRIM therapeutically lies in avoiding its potential to induce or exacerbate autoimmune pathology. Here, we describe a unique form of TRIM in AMs, driven by WTAP-mediated m^6^A hypomethylation, which enhances antimicrobial phagocytosis without activating pro-inflammatory genes. Distinguished by this lack of hyperresponsiveness, this epitranscriptomic program contrasts with conventional training induced by stimuli like β-glucan (23), IL-1β (22), or the bacillus Calmette–Guérin (BCG) vaccine (74). This ‘balanced’ reprogramming is particularly advantageous, as it suggests a potential pathway to harness the protective benefits of TRIM while mitigating the risk of exacerbating inflammatory or autoimmune pathology, which is a significant concern in therapeutic immune training.

In conclusion, we report a previously undiscovered epitranscriptomic mark involved in TRIM establishment in AMs. This innate immune memory driven by WTAP-mediated m^6^A hypomethylation integrated with metabolic reprogramming confers prolonged protection against secondary bacterial pneumonia. Hence, this work not only advances our fundamental understanding of epitranscriptomic control in innate immune memory but also opens new avenues for therapeutic intervention against secondary bacterial infections and possibly other macrophage-driven chronic lung diseases.

## Methods

### Study approval and biosafety statement

All procedures and methods were conducted in accordance with regulations of Sun Yat-sen University and Guangzhou National Laboratory. Animal experiments were approved by the Institutional Animal Care and Use Committee (IACUC) of Sun Yat-sen University (No. SYSU-IACUC-2021-B1310). All the mouse experiments involving H1N1 infection were approved and performed according to a protocol approved by the IACUC of the Institute of Guangzhou National Laboratory (No. GZLAB-AUCP-2025-06-A05). All cell experiments with viruses or bacteria were conducted in the biosafety level 2 laboratory (BSL-2) of the Guangzhou National Laboratory. All experiments strictly followed standard operating procedures, and all waste materials were autoclaved and verified before disposal.

### Mice

C57BL/6 mice were purchased from Sun Yat-sen University Laboratory Animal Center (Guangzhou, China). C57BL/6 *Lyz*M-Cre^+^ *Wtap*^-/-^ mice were generated by GemPharmatech Co., Ltd. using the CRISPR/Cas9 approach (38). Mice were housed in a specific pathogen-free (SPF) facility with ad libitum access to food and water and were maintained on a 12-h light/12-h dark cycle, with 50–60% humidity and at 20–25 °C at Sun Yat-sen University Laboratory Animal Center. All experiments were performed in accordance with the guidelines from the Institutional Animal Care and Use Committee (IACUC) of Sun Yat-sen University.

### Bacterium and virus strains

*Pseudomonas aeruginosa* (*P. aeruginosa*, ATCC27853), *Escherichia coli* (*E. coli*, ATCC11303) and Enhanced Yellow Fluorescent Protein expressing *Escherichia coli* (eYFP-*E. coli*) were grown in LB broth under constant shaking at 37 °C overnight.

Influenza A virus (strain A/Puerto Rico/8/1934, H1N1) were generously gifted from Dr. Nan Qi (Guangzhou National Laboratory). For viral amplification, IAV was propagated in 10-day-old SPF embryonated chicken eggs via the allantoic route. For plaque assays, MDCK cells were seeded in 6-well plates and infected with serially diluted virus supernatants at 37 °C for 1 h. The cells were then washed with PBS and overlaid with DMEM (Gibco) containing 2% FBS, 2 μg/mL TPCK-trypsin (Sigma) and 2% SeaPlaque agarose (Lonza) at 4 °C for 30 min. The plates were incubated upside down at 37 °C for a further 48 h, followed by counting visible plaques for viral titer determination.

### Materials

Information about reagents and resources is listed in Supplementary Table 1.

### Isolation of AM cells from pulmonary alveoli

Bronchoalveolar lavage (BAL) was performed and BAL fluid (BALF) was collected as previously described with minor modifications (35). In brief, place the euthanized mice in supine position and make an incision at the cervical region using surgical scissors and removing the skin. After exposure and midline incision of the trachea, make a small horizontal incision on the trachea without severing. Slowly insert an 18-gauge cannula into the trachea, with the needlepoint facing toward the lungs. Secure the cannula in place by tying a surgical thread around the trachea. Mouse airways were flushed with a total of 5 mL endotoxin-free saline by instillation of 0.5 mL aliquots. After collecting the BALF from all the experimental mice, centrifuge the BALF at 2000 × g for 5 min at 4 °C. Resuspend the cell pellets in 200 μL of PBS and keep at 4 °C until use.

### Challenge of mice

For experiments with influenza model, isoflurane-anesthetized mice were challenged with intranasal (i.n.) IAV infection (150 PFU in 20 μL of sterile 1 × PBS), or 1 × PBS only (controls). For experiments with pulmonary inflammation, isoflurane-anesthetized mice were administered with intratracheal (i.t.) LPS (10 mg/Kg, *E. Coli* O111:B4, Sigma; reconstituted in sterile 1 × PBS), or 1 × PBS only (controls).

For experiments with lung bacterial infection, isoflurane-anesthetized mice were challenged with intratracheal (i.t.) *P. aeruginosa* infection (ATCC27853, 5 × 10^7^ CFU), or 1 × PBS only (controls). For experiments with STC-15 pretreatment, isoflurane-anesthetized mice were challenged with intratracheal (i.t.) STC-15 (10 mg/kg, Selleck; dissolved in the adjuvant composed of 5% DMSO, 40% PEG300, 5% Tween80 and 50% ddH_2_O), or adjuvant only (controls).

### Estimation of Bacterial Burden *In Vivo*

Mice were infected (i.v.) with *P. aeruginosa* (1.5 × 10^7^ CFU per mouse). After 12 h of infection, mice were euthanized. The BALF was collected and centrifuged at 2000 × g for 5 min at 4 °C. 10 µL supernatant fluids were diluted with 20 µL sterile PBS and then plated on BHI Agar plates. After 24 h incubation at 37 °C, the bacterial colonies were enumerated.

### Phagocytosis assays

AMs were isolated by BALF, seeded at equal numbers and left to adhere in RPMI medium (10% FBS, 1% PS, 100 ng/mL GM-CSF). After 12 h, cells were washed with 1 × PBS and incubated with eYFP-*E. coli* (2.5 × 10^6^ CFU) for 1 h. Subsequently, cells were washed again and treated with proteinase K (50 µg/mL; Roche) for 10 min at 4°C to remove residual adherent bacteria. After incubation with indicated times, cells were digested with Trypsin, washed with 1 × PBS, and then analyzed by flow cytometry to quantify eYFP⁺ cells.

### Chemotaxis assays

Chemotaxis assay was performed using Transwell insert (Corning) with 8 μm pores following a previously described procedure with minor modifications (12). Briefly, 1×10^5^ AMs were seeded into the top chamber, while the lower chamber contained RPMI 1640 medium supplemented with 5% FBS, 5% fresh mouse serum (to induce C5a production) and 1 × 10^7^ CFU *P. aeruginosa*. Cells were allowed to migrate for 1 or 2 hours in 37 °C incubator with 5% CO2. After migration, the inserts were stained with crystal violet stain (Beyotime), Non-migrated cells on the upper side of the membrane were carefully removed using a cotton swab. The transwell membranes were then removed and mounted on slides. Cells from five random high-power fields (HPF) per membrane were imaged using an inverted microscope equipped with a camera, manually counted, and the total number of migrated cells per insert was reported as the mean of three technical replicates.

### Cathepsin activity assays

Lysosomal cathepsin B (CTSB) enzymatic activity was detected using Cathepsin B Assay (Magic Red) reagents (Abcam) according to the manufacturer’s protocol. Cells cultured on confocal dish were incubated with Magic Red CTSB reagents for 15 min at 37 °C, followed by staining with Hoechst 33342. Images were acquired using confocal microscopy.

### Seahorse assays

Real-time cell metabolism of AMs was measured by using the Seahorse XF Cell Mito stress test kit and the Seahorse XF glycolytic stress test kit (Agilent Technologies) according to the manufacturer’s instructions. Briefly, purified AMs were seeded into XF 96-well plates (Agilent Technologies) at a density of 50,000 cells per well and cultured overnight in complete RPMI 1640 medium. Prior to the assay, cells were washed and equilibrated for 1 h in Seahorse assay medium (Agilent Technologies) supplemented with either 2 mM glutamine (glycolytic stress test) or an additional 10 mM glucose and 1 mM pyruvate (mito stress test). The following compounds were applied at the indicated final concentrations: 1.5 μM oligomycin, 4 μM FCCP and 0.5 μM rotenone/antimycin A in the mito stress test or 10 mM glucose, 1 μM oligomycin and 50 mM 2-DG in the glycolytic stress test. Oxygen consumption rate (OCR) and extracellular acidification rate (ECAR) were measured using a Seahorse XFe96 analyzer (Agilent Technologies), and data were analyzed using Wave Desktop software (v2.6).

### Lung macroscopic analysis and histopathology

The indicated mice were euthanized by CO2 exposure. Then, the lungs were removed and fixed in 4% paraformaldehyde (MA0192, Meilunbio) for more than 24 h and embedded in paraffin. Haematoxylin & eosin (H&E) staining of the lung sections (thickness, 6 μm) was performed by Servicebio of China. The H&E-stained sections were examined by microscopy (DMi8, Leica). All histology analyses were conducted in a blinded manner.

### Flow cytometry

Leukocyte populations in the BALF were identified and quantified by flow cytometry. In brief, BALF was collected from mice by lavaging the lungs. The BALF was centrifuged at 1000 × g for 10 min at 4 °C. The cell pellet was resuspended in 500 μL of red blood cell lysis buffer (Beyotime, C3702) and incubated for 5 min at room temperature. After centrifugation, cells were washed twice with PBS.

For surface marker staining, cells were incubated with Fc receptor blocking antibody (anti-CD16/32) for 10 min at 4 °C, followed by staining with fluorochrome-conjugated specific antibodies for 30 min at 4 °C in the dark. Cells were then washed twice with PBS, passed through a 40 μm cell strainer (BD Falcon), and resuspended in 200 μL of PBS for acquisition.

Flow cytometry was performed on a CytoFLEX cytometer (Beckman), and data were analyzed using FlowJo software (Flowjo V10). Alveolar macrophages were defined as CD45⁺CD11c⁺Siglec-F⁺ cells, and neutrophils as CD45⁺CD11b⁺Ly6G⁺ cells.

### Immunofluorescence staining and confocal microscopy

For Immunofluorescence staining, AMs were seeded at concentration of 6×10^4^ per well in a confocal dish. After being washed with 1×PBS three times, cells were fixed with 4% paraformaldehyde (Meilunbio) for 15 min, then permeabilized using PBS containing 0.1% Triton X-100 for 10 minutes and blocked for 1 h in blocking buffer at room temperature. Samples were incubated with the indicated primary antibodies overnight at 4 ℃, then washed in PBS three times and incubated with the corresponding fluorescent secondary antibody for 1 h at room temperature, followed by washing with PBS three times. Samples were then counterstained with DAPI and imaged with Spinning disk confocal microscope (Nikon ECLIPSE Ti2-E+CSU-W1).

### Live-cell imaging

For live imaging of cellular movement of AMs, AMs were seeded at concentration of 4×10^4^ per well in a confocal dish. Three hours prior to imaging, AM cells were stained with PKH26 (Sigma‒Aldrich) at a final concentration of 2 μM per dish. In order to allow robust identification of nuclei, the cultures were incubated with the membrane-permeable dyes Hoechst 33342 (Beyotime) for 10 min at 37 ℃. The cells were washed three times with PBS prior to imaging. Then, confocal imaging was performed on a Spinning disk confocal microscope (Nikon ECLIPSE Ti2-E+CSU-W1) at 37 °C and 5% CO2 in growth medium. An oil immersion objective (100×) was used to acquire high-resolution images. The motorized stage was programmed to visit multiple locations in different wells with intervals of 1 or 1.5 minutes per location, and each imaging session lasted for 1 hour. Images were exported as nd2 file format for further analysis in NIS - elements Viewer. The positional data of cells were exported to Excel files and subsequently analyzed for statistical imaging using R (4.4.1).

### Preparation of LNP

The lipid nanoparticles (LNPs) used in this study were formulated and loaded with siRNA by GenScript (Nanjing, China) using a standardized microfluidic mixing protocol. Briefly, an aqueous phase containing siRNA in citrate buffer (pH 4.0) and an ethanol phase containing the lipid mixture (SM-102, cholesterol, DMG-PEG2000, DSPC, and DSPE-PEG2k-Mannose) were combined at a 3:1 volume ratio in a microfluidic device to instantaneously form mRNA-loaded LNPs. The particles were further purified through overnight dialysis against 1× PBS at 4°C to remove residual ethanol. Finally, the LNP concentration was adjusted via ultrafiltration and sterilized by passage through a 0.2-μm membrane filter.

The physicochemical properties of the LNPs were characterized through multiple analytical methods. Particle size, polydispersity index (PDI), and zeta potential were determined by dynamic light scattering (DLS) using a Zetasizer PRO (Malvern). RNA encapsulation efficiency was quantified with a Quant-iT RiboGreen RNA assay (Thermo) according to the manufacturer’s instructions.

### Liquid chromatography–tandem mass spectrometry (LC–MS/MS) assays

LC–MS/MS assays were performed as previously described (38).

### RNA decay assays

AM cells with KO of *Wtap* and control or treatment with STC-15 were treated with Act D (5 μg/mL; Sigma) for 0, 2, 4, 6, 8, 10 and 12 h, respectively, and then RNA samples were extracted for qRT‒PCR to determine the mRNA levels of the indicated genes. Specifically, total RNA from cells was extracted using TRIzol (Invitrogen). RNA quantity and quality were assessed with NanoDrop 2000 spectrophotometer (Thermo Fisher Scientific). Then, 1 µg of total RNA used for cDNA synthesis with *Evo M-MLV* RT Mix Kit with gDNA Clean for qPCR (AG), and the diluted cDNA was used as template for qPCR with 2×Polarsignal^®^ qPCR Mix (MIKX). Target mRNA expression levels were normalized to the expression level of *Actb* or *Gapdh* in each individual sample. The 2^−ΔΔCt^ method was used to calculate relative expression changes. Specific primers were designed for each gene transcript and are listed in Supplementary Table 2.

### m^6^A RNA-IP-qRT–PCR (MeRIP‒qPCR)

MeRIP assay is conducted using the EpiQuik™ CUT&RUN m^6^A RNA Enrichment (MeRIP) Kit (EPIGENTEK) according to the manufacturer’s instructions with minor modifications. Briefly, total RNA (10 µg) was immunoprecipitated with a certain amount of RNA (1/10) reserved as the input control (Input). Immunocapture involved mRNA samples, m^6^A antibodies, affinity beads and buffer vortexed for 90 min at room temperature. RNA fragmentation was achieved using cleavage enzymes, followed by proteinase K (Roche) treatment and RNA purification to isolate m^6^A-enriched RNA. The immunoprecipitated m^6^A RNA (IP) was then subjected to RT-qPCR analysis, with primers provided in Supplementary Table 2.

### RNA-seq and data analysis

Whole-cell total RNA was isolated using TRIzol reagent (Invitrogen) and quantified using a NanoDrop 2000 spectrophotometer (Thermo Fisher Scientific). The cDNA library was constructed by Biomarker Technologies. Sequencing was performed on an Illumina HiSeq 2500 platform. High-quality reads were mapped to the mouse reference genome (mm9) using HISAT2. DESeq, an R package, was applied for differential gene expression analysis. We filtered the differentially expressed genes based on a false discovery rate (FDR) <0.05.

### Direct RNA sequencing (ONT DRS) and data analysis

1. **Nanopore direct RNA sequencing**. Nanopore direct RNA sequencing (ONT DRS) was conducted by Benagen Technology Co., Ltd (Wuhan, China). The total RNA from WT and *Wtap*-cKO AMs was extracted using TRIzol (Invitrogen), and the quality and quantity of RNAs were assessed using the NanoDrop 2000 spectrophotometer (Thermo Fisher Scientific). Each total RNA sample (50 μg) was subjected to mRNA enrichment using the mRNA Dynabeads™ Kit (Thermo Fisher Scientific). The mRNA was then ligated to the Nanopore RT Adapter using T4 DNA ligase (NEB). Following reverse transcription of this adapter-ligated mRNA, the cDNA product was purified with Agencourt RNAClean XP beads. Subsequently, the RNA adapter was added, and the ligation product was purified with Agencourt RNAClean XP beads (Beckman) once more. Finally, the prepared library was loaded onto an R9.4 sequencing chip and sequenced on a PromethION device (Oxford Nanopore Technologies) for 48 to 72 hours.
2. **Nanopore basecalling and m^6^A detection**. Raw electrical signal data (POD5 format) were processed with Dorado (v1.2.0, Oxford Nanopore Technologies). First, basecalling was performed using the High Accuracy (HAC) model (rna004_130bps_hac@v5.2.0). Subsequently, m^6^A modifications were detected concurrently using the dedicated modified base model (rna004_130bps_hac@v5.2.0_m6A_DRACH@v1). During processing, reads were aligned to the GRCm38.p6 reference genome via Dorado’s integrated Minimap2 aligner (using the --reference argument), which also enabled the direct output of aligned BAM files containing modification probability tags (MM/ML). To generate site-level modification profiles, aligned BAM files were processed with Modkit (v0.5.0). The pileup command was used to aggregate modification probabilities, calculating m^6^A frequency and coverage per adenine site, with the results output in bedMethyl format.
3. **m^6^A data pre-processing and differential analysis**. Differential m^6^A analysis was performed with the methylKit R package (v1.34.0). Modification percentages from Modkit were reformatted into m^6^A (freqC) and unmodified adenine (freqT) frequencies for compatibility and imported via methRead. Following data import, quality control involved removing low-coverage sites (< 200 reads) with filterByCoverage and normalizing coverage using normalizeCoverage. Samples were merged with the unite function, and Fisher’s exact test (calculateDiffMeth) was applied to define differential sites (|meth.diff| ≥ 25%, p < 0.01).

### Statistical analysis

The statistical analysis was done using Microsoft Excel software and GraphPad Prism to assess the differences between experimental groups. Statistical significance was determined using a two-tailed, unpaired Student’s *t*-test between two groups. P ≤ 0.05 was considered to indicate a statistically significant difference. All experiments were performed three or more times independently under identical or similar conditions.

## Data and code availability

Publicly available datasets analyzed in this study including GSE222150, GSE208294, GSE145926 and GSE136831 are available at the GEO database. Bulk RNA-Seq and Direct RNA-Seq raw data generated for this study have been deposited in the Sequence Read Archive (SRA) under accession numbers PRJNA1401683 and PRJNA1403425, respectively.

## Acknowledgments

This work was supported by projects from the National Natural Science Foundation of China (32571027, U23A6012 and 32200712), the Guangdong Science and Technology Department (2024B1515040009, 2025B0303000010 and 2023B1212060028), the Fundamental Research Funds for the Central Universities (23yxqntd001), the Sun Yat-sen University–ImmunoArt Biotechnology Company Joint Research Center for Autoimmune Disease (HT-99982024-0483), and the Major Project of Guangzhou National Laboratory (GZNL2024A01016). The authors thank the technical assistance of Hongmei Li with electron microscopy, Haonan Yu with confocal microscopy, Yingxia Wu with flow cytometry, and Xin Cai with LC–MS/MS.

## Author contributions

Y.G., A.X., and S.Y. conceived the study, analyzed the data, and prepared and wrote the manuscript with input from the other authors. Y.G., X.H., Y.C., R.C., H.W., Z.Q., W.S., and J.H. performed the experiments collaboratively. Y.G., X.H., and Z.L. performed the data analysis of RNA-seq, scRNA-seq, and ONT DRS. Y. Z. and N.Q. provided key technical guidance for several experimental procedures. S.Y. and A.X. led the project, supervised and coordinated the research. All authors have read and approved the manuscript.

