## Supplementary materials for "WTAP-mediated epitranscriptomic program in alveolar macrophages confers prolonged protection against postinfluenza bacterial pneumonia"

### Supplemental figures and figure legends

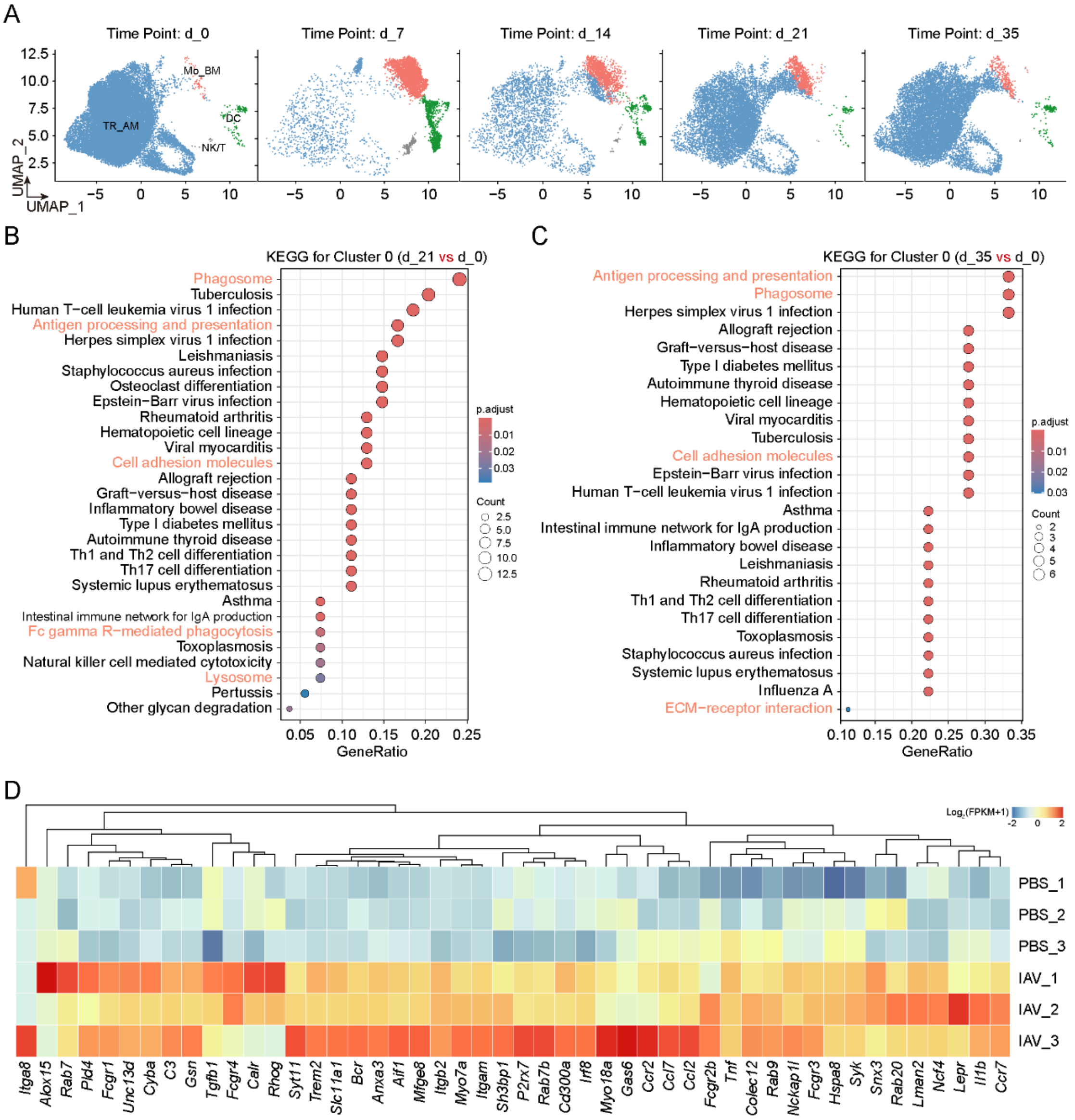


**Supplemental Figure 1. Respiratory virus-trained AMs are characterized by a high phagocytic capacity.** (**A**) UMAP depiction of BALF cell clusters at the respective time points p.i. (**B** and **C**) KEGG enrichment of upregulated genes in cluster 0 from IAV-infected mice on day 21 (B) or day 35 (C) after infection versus those from uninfected mice. (**D**) Heat maps showing the abundance of different gene transcripts related to phagocytosis in AMs from PBS- or IAV-infected mice.

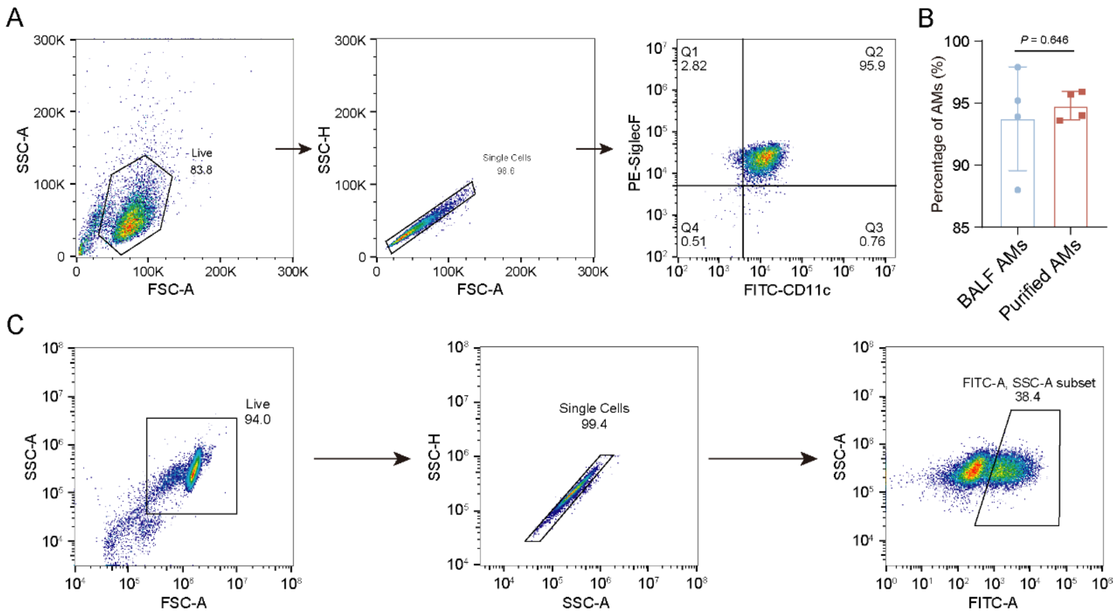


**Supplemental Figure 2. The gating strategy of flow cytometry.** (**A**) The gating strategy for identifying extracted and purified AMs. **(B)** The proportion of AMs in BALF and after purification. (**C**) The gating strategy for AMs that phagocytose eYFP-*E. coli*.

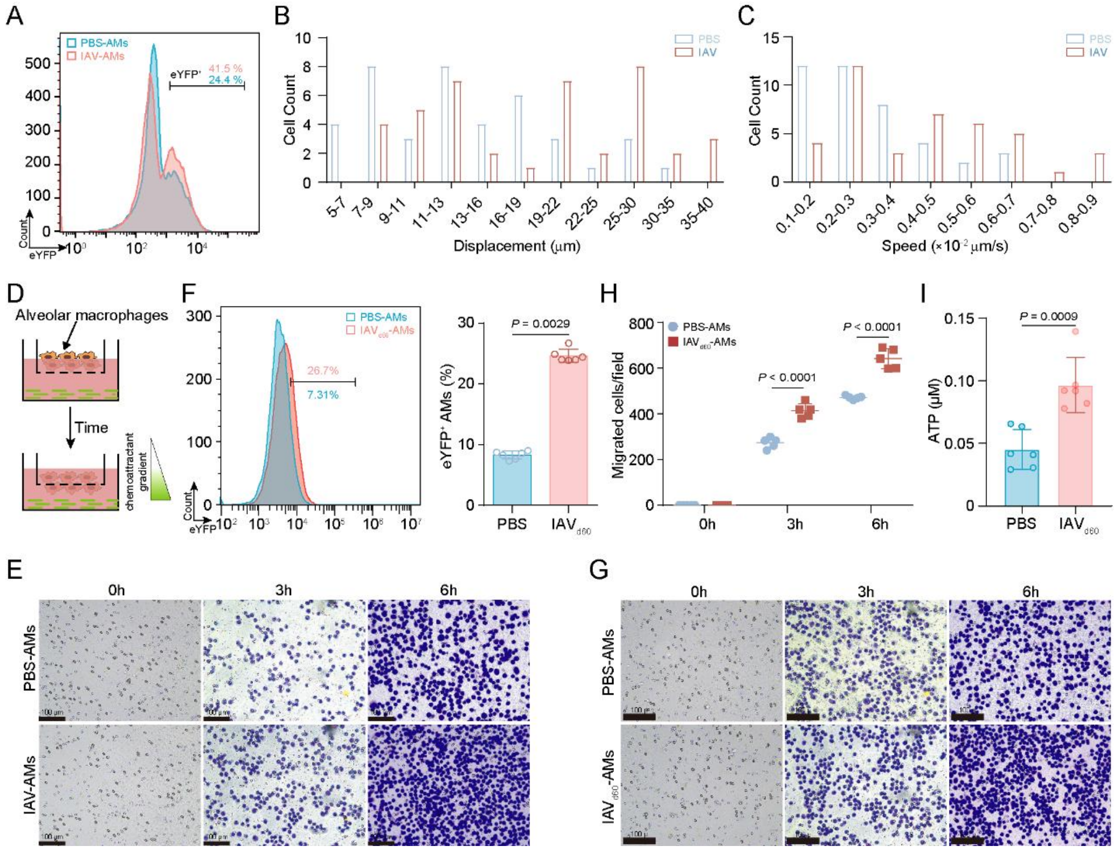


**Supplemental Figure 3. IAV-trained AMs have enhanced phagocytic and hydrolytic functions.** (**A**) Representative flow cytometry detection of YFP$^+$ AMs after co-incubation of PBS- or IAV-AMs with eYFP-*E. coli*. (**B** and **C**) Histogram of displacement in micrometers (B) and average track velocity (micrometers per second) (C) of AMs from control or IAV-infected mice during a 1 h intravital imaging session. (**D**) Cartoon showing the *in vitro* chemotaxis assay, and *P. aeruginosa* was used as a stimulus for AM migration. (**E**) Representative imaging of migrated AMs after crystal violet staining. Scale bar, 100 μm. (**F**) Representative flow cytometry detection of eYFP$^+$ AMs after co-incubation of PBS- or $IAV_{d60}$-AMs with eYFP-*E. coli* (left) and quantifying the phagocytosis of PBS- and IAV-AMs on eYFP-*E. coli* (right). n = 8 mice per group. (**G** and **H**) Representative imaging of migrated AMs after crystal violet staining (G) and quantification of the *in vitro* chemotaxis assay of PBS- and $IAV_{d60}$-AMs (H). Scale bar, 100 μm. n = 5 mice per group. (**I**) Determination of ATP concentration in PBS- and IAVd60-AMs. n = 6 mice per group.

Data are presented as the mean ± s.d. in F, H and I with individual measurements overlaid as dots, and statistical analysis was performed in F and I using a two-tailed Student's *t*-test or in H using Multiple Comparisons Following Two-Way ANOVA. Data in A and E-G are representative of three independent biological experiments.

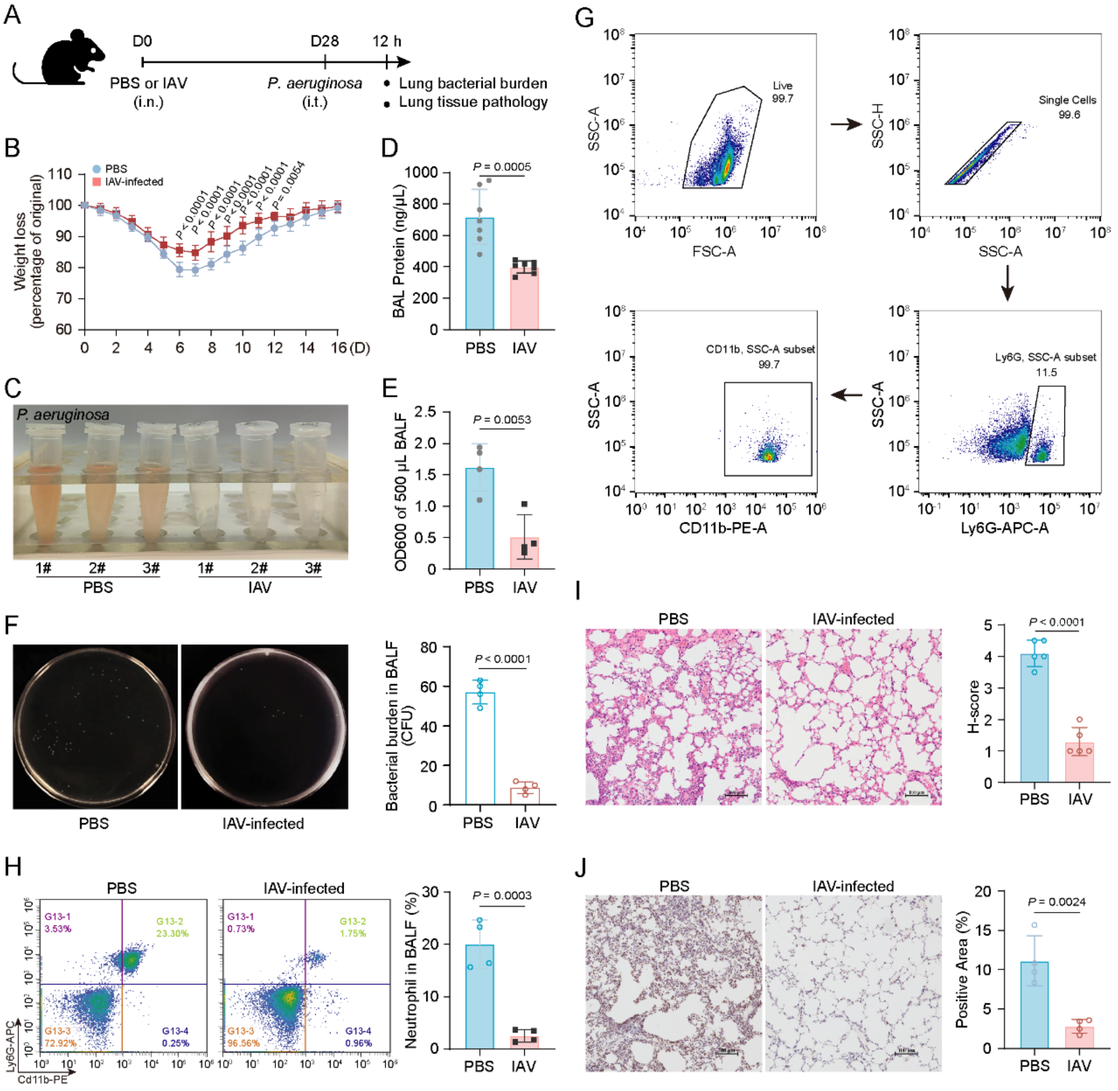


**Supplemental Figure 4. IAV-trained AMs protects mice against pulmonary bacterial infection.** (**A**) Experimental schema of i.n. IAV infection in mice followed by i.t. *P. aeruginosa* infection. (**B**) Weight loss monitored over time of control or IAV-infected mice infected with *P. aeruginosa*. n = 8 mice per group. (**C** to **E**) Observation of the bleeding in BALFs (C) and measurement of the BALF protein content (D) and turbidity of BALFs (E) from adjuvant- and STC-15-pretreated mice 12 hours after *P. aeruginosa* administration. n = 3, 7, or 4 mice per group. (**F**) Representative bacterial plate pictures of BALFs from control or IAV-infected mice at 12 h after *P. aeruginosa* infection (left) and quantification of bacterial burden (right). n = 4 mice per group. (**G**) The gating strategy for neutrophils. (**H**) Flow cytometric plot shows Cd11b$^{+}$ Ly6G$^{+}$ neutrophils in BALFs from control or IAV-infected mice at 12 h after *P. aeruginosa* infection (left) and quantification of the percentage of neutrophils (right). n = 4 mice per group. (**I**) Representative histology images of lung tissues from control or IAV-infected mice at 12 h after *P. aeruginosa* infection (left) and histochemistry score of

H&E staining (right). Scale bar, 100 μm. n = 5 mice per group. (**J**) Representative histology images of lung tissues from control or IAV-infected mice at 12 h after *P. aeruginosa* infection (left) and quantification of the percentage of MPO staining (right). Scale bar, 100 μm. n = 4 mice per group. Data are presented as the mean ± s.d. in D-F and H-J with individual measurements overlaid as dots, and statistical analysis was performed in D-F and H-J using a two-tailed Student's *t*-test or performed in B using Multiple Comparisons Following Two-Way ANOVA. Data in C, F and H-J are representative of three independent biological experiments.

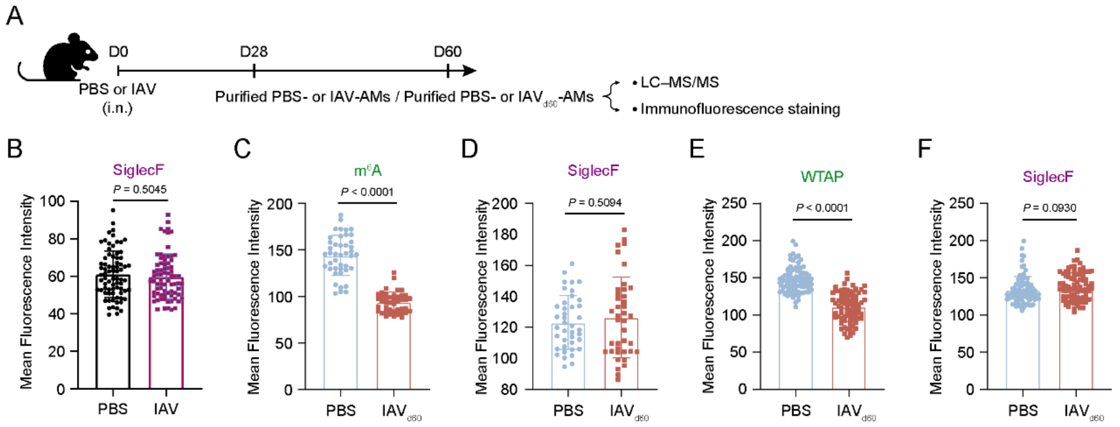


**Supplemental Figure 5. WTAP is downregulated in AMs during IAV-induced trained immunity.** (**A**) Experimental schema of i.n. IAV infection in mice and subsequent phenotypic detections. (**B**) Quantification of Siglec F intensity in PBS- and IAV-AMs. (**C** and **D**) Quantification of $m^6A$ (C) and Siglec F (D) immunofluorescence intensity in AMs from PBS- and day 60 IAV-infected ($IAV_{d60}$) mice. (**E** and **F**) Quantification of WTAP (E) and Siglec F (F) immunofluorescence intensity in AMs from PBS- and day 60 IAV-infected ($IAV_{d60}$) mice.

Data are presented as the mean ± s.d. in B-F with individual measurements overlaid as dots, and statistical analysis was performed using a two-tailed Student's *t*-test.

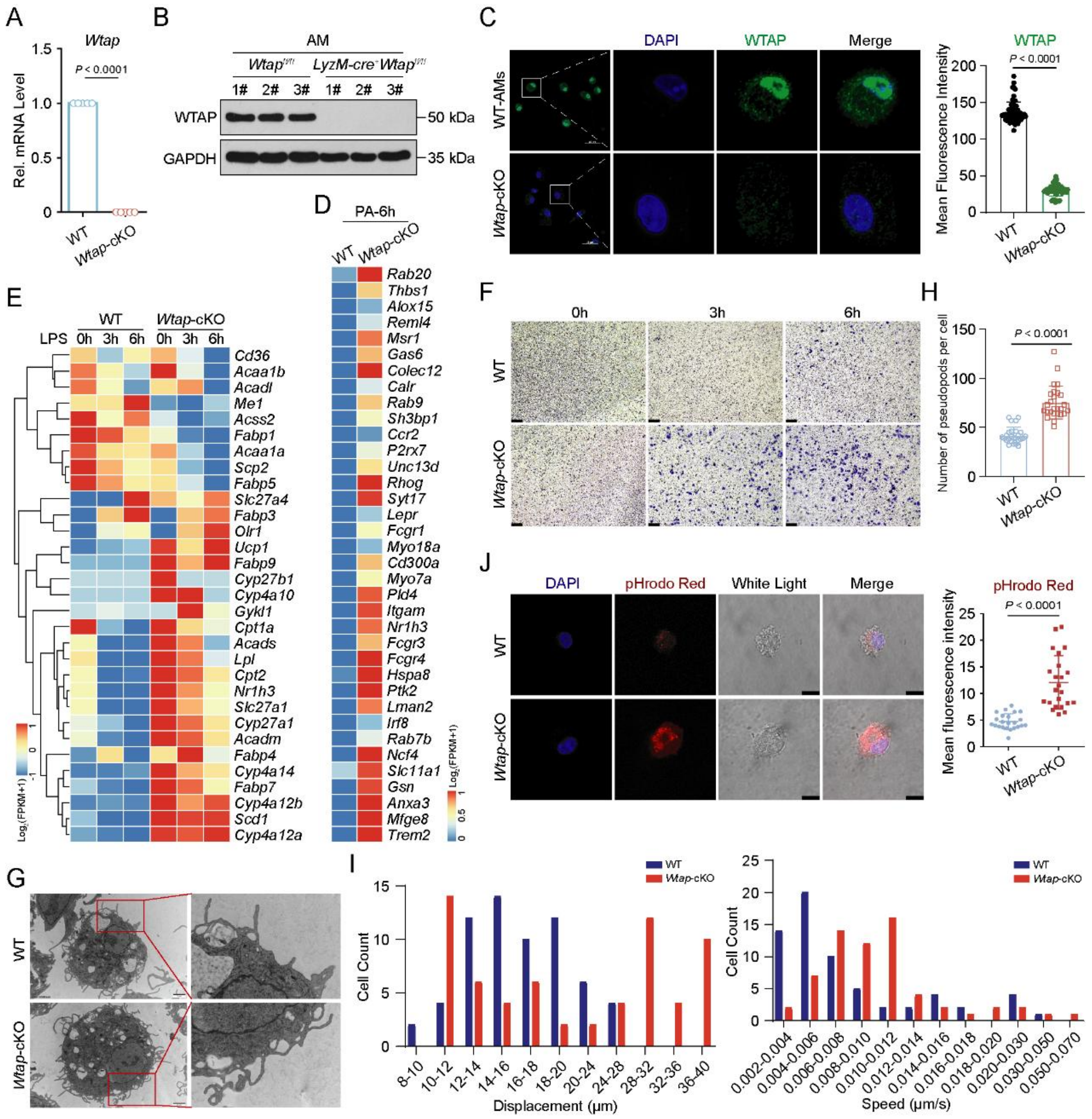


**Supplemental Figure 6. WTAP-deficient AMs exhibit enhanced phagocytic activity.** (**A** and **B**) qRT-PCR (A) and Immunoblotting (B) analysis of WTAP expression in WT and *Wtap*-cKO AMs. n = 3 or 5 mice per group. (**C**) Representative imaging of WTAP immunofluorescence staining in WT and *Wtap*-cKO AMs (left) and quantification of WTAP intensity (right). Scale bar, 25 μm. (**D**) RNA-seq heat maps showing the mRNA abundance of genes related to phagocytosis in WT and *Wtap*-cKO AMs infected with *P. aeruginosa* (PA). (**E**) RNA-seq heat maps showing the mRNA abundance of genes related to the PPARγ signaling pathway. (**F**) Representative imaging of migrated WT and *Wtap*-cKO AMs after crystal violet staining. Scale bar, 100 μm. (**G** and **H**) Representative TEM micrograph of pseudopodia in WT and Wtap-cKO AMs (G) and quantification of pseudopod numbers (H). (**I**) Histogram of displacement in micrometers (left) and average track velocity (micrometers per second) (right) of WT and *Wtap*-cKO AMs during a 1 h intravital imaging session.

(**J**) Representative imaging of pHrodo Red in WT and *Wtap*-cKO AMs (left) and quantification of fluorescence intensity (right). Scale bar, 25 μm.

Data are presented as the mean ± s.d. in A, C, H and J with individual measurements overlaid as dots, and statistical analysis was performed using a two-tailed Student's *t*-test. Data in B, C, F, G and J are representative of three independent biological experiments.

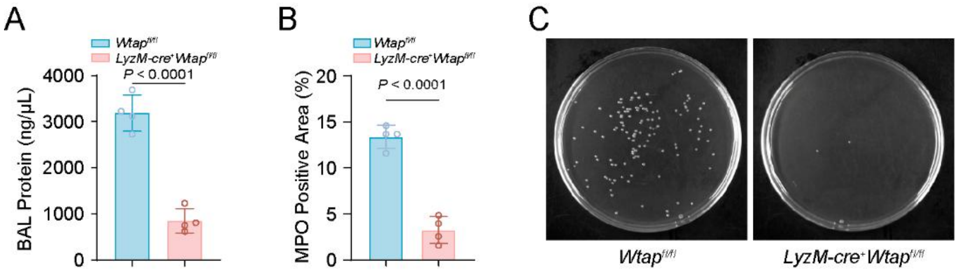


**Supplemental Figure 7. WTAP deficiency in AMs protects mice against pulmonary bacterial infection.** (**A**) Quantification of BAL protein from *Wtap*$^{fl/fl}$ and *LyzM*-Cre$^{+}$ *Wtap*$^{fl/fl}$ mice 12 hours after *P. aeruginosa* administration. n = 4 mice per group. (**B**) MPO staining of lung tissues from *Wtap*$^{fl/fl}$ and *LyzM*-Cre$^{+}$ *Wtap*$^{fl/fl}$ mice at 12 h after *P. aeruginosa* infection. (**C**) Representative bacterial plate pictures of BALFs from *Wtap*$^{fl/fl}$ and *LyzM*-Cre$^{+}$ *Wtap*$^{fl/fl}$ mice at 12 h after *P. aeruginosa* infection.

Data are presented as the mean ± s.d. in A and B with individual measurements overlaid as dots, and statistical analysis was performed using a two-tailed Student's *t*-test. Data in C are representative of three independent biological experiments.

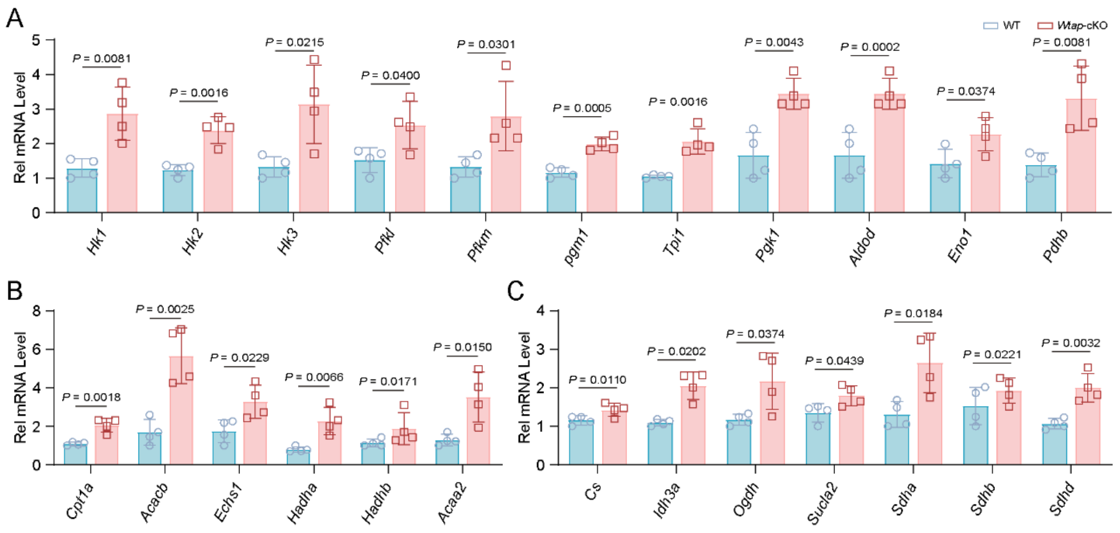


**Supplemental Figure 8. Metabolic rewiring in WTAP-deficient AMs enhance the characteristics of trained immunity.** (**A** to **C**) qRT-PCR showing the mRNA abundance of enzyme genes involved in glycolysis (A), fatty acid oxidation (B) and TCA cycle (C) in WT and *Wtap*-cKO AMs.

Data are presented as the mean ± s.d. in A-C with individual measurements overlaid as dots, and statistical analysis was performed using a two-tailed Student's *t*-test.

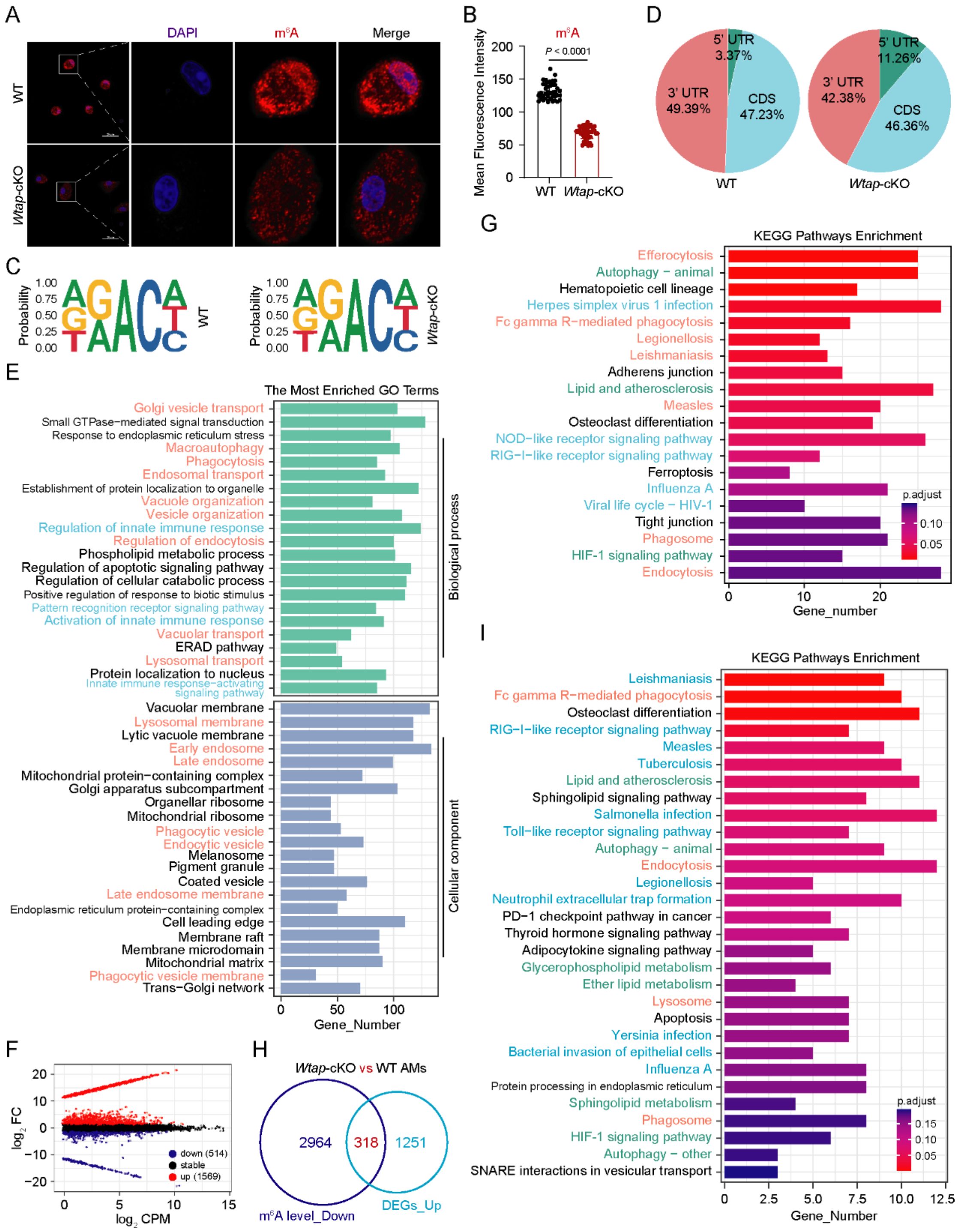


**Supplemental Figure 9. WTAP regulates AM trained immunity through m$^6$A modification**. (**A** and **B**) Representative imaging of m$^6$A (red) immunofluorescence staining in WT and *Wtap*-cKO AMs (A) and quantification of m$^6$A intensity (B). Scale bar, 25 μm. (**C**) Predominant consensus motif "RRACH" was detected in WT and *Wtap*-cKO AMs. (**D**) Pie charts depicting the proportion of m$^6$A peak distribution in the 5' UTR, CDS, and 3' UTR regions across mRNA transcriptome. (**E**) GO enrichment analysis of the genes with decreased m$^6$A marks in *Wtap*-cKO AMs compared with

WT controls. (**F**) Volcano plots showing the differentially expressed genes in WT and *Wtap*-cKO AMs. (**G**) KEGG enrichment analysis of the upregulated genes in *Wtap*-cKO AMs compared with WT controls. (**H**) Venn diagrams showing transcripts with decreased $m^6A$ abundance and upregulated mRNA expression in *Wtap*-cKO AMs compared with WT controls. (**I**) KEGG enrichment analysis of the overlapping genes presented in (H).

Data are presented as the mean ± s.d. in B with individual measurements overlaid as dots, and statistical analysis was performed using a two-tailed Student's *t*-test. Data in A are representative of three independent biological experiments.

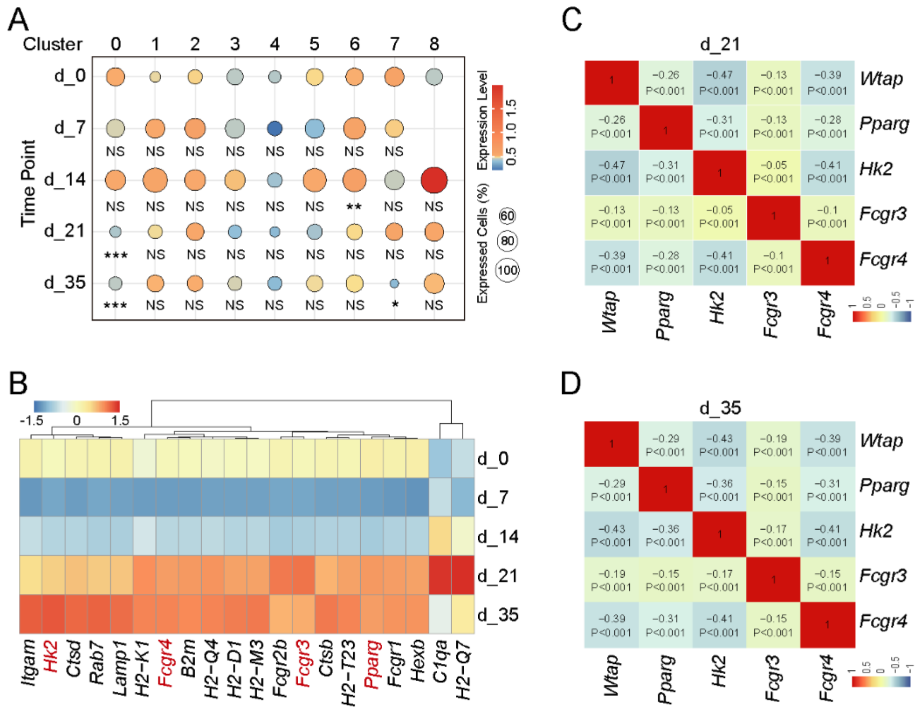


**Supplemental Figure 10. The downregulation of *Wtap* is associated with a significant upregulation of phagocytic genes.** (**A**) Dot plot graph showing expression of WTAP in each indicated cluster from all time points. (**B**) Heatmap showing expression of the genes involved in phagocytosis, antigen processing and presentation and metabolism in cluster 0 from all IAV-infected time points. (**C** and **D**) Spearman correlations between the expression of *Wtap* and its target gene (*Hk2*, *Pparg*, *Fcgr3* and *Fcgr4*) in cluster 0 from IAV-infected mice on day 21 (C) and day 35 (D). Statistical analysis was performed in A using Mann-Whitney U test.

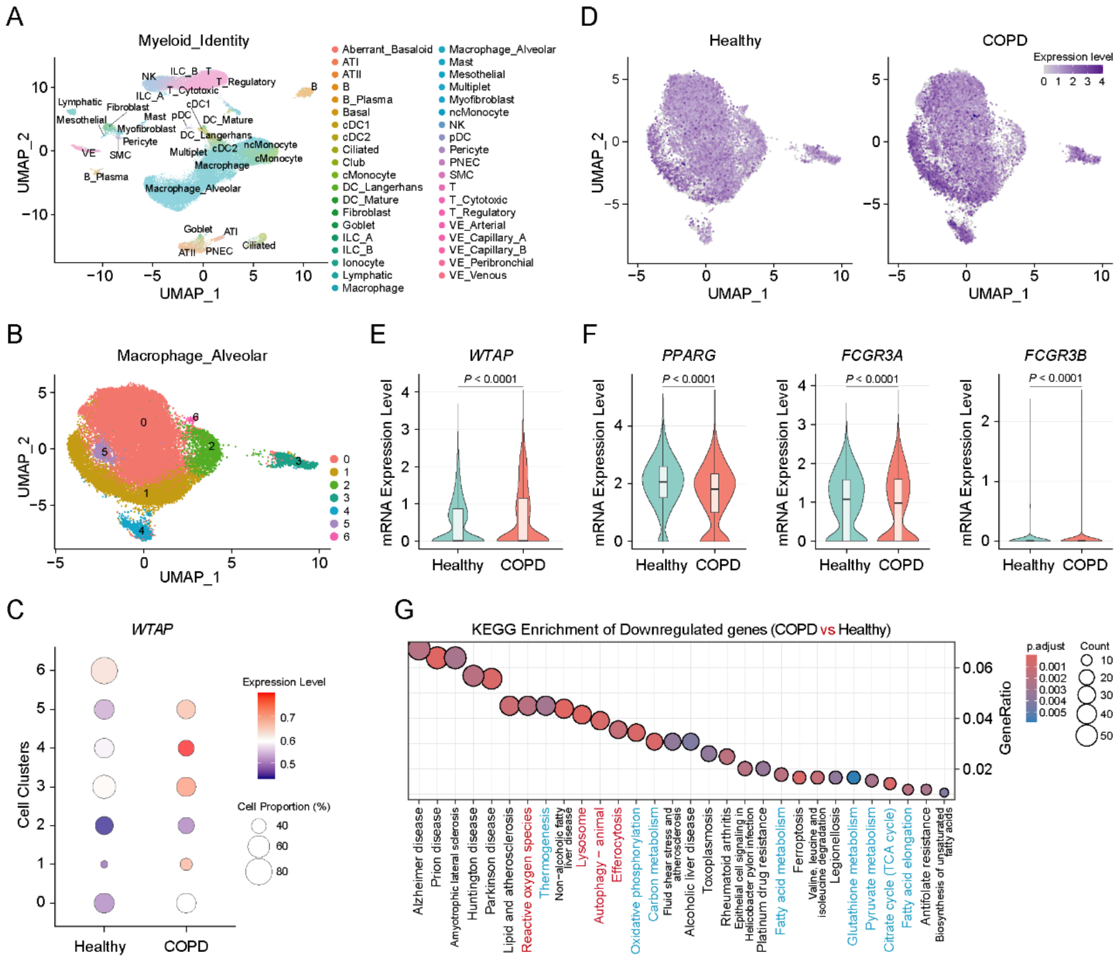

**Supplemental Figure 11. Trained AMs associated with lower expressed WTAP alleviate the severity of pulmonary diseases.** (**A**) UMAP representation of major cell types from 18 COPD, and 28 control donor lungs; each dot represents a single cell, and cells are labeled as one of 39 discrete cell varieties. AT, alveolar type; cDC, classical dendritic cell; pDC, plasmacytoid dendritic cell; NK, natural killer; ILC, innate lymphoid cell; PNEC, pulmonary neuroendocrine cell; SMC, smooth muscle cell; VE, vascular endothelial. (**B**) UMAP presentation of alveolar macrophages and associated clusters. (**C**) Dot plot graph showing expression of *WTAP* in each indicated cluster from control donors and COPD patients. (**D**) UMAP projection of the expression of *WTAP* in alveolar macrophages from healthy donors and COPD patients. (**E** and **F**) Violin plot showing the mRNA expression of *WTAP* (E) or *PPARG*, *HK2* and *FCGR3A* (F) in alveolar macrophages from healthy donors and COPD patients. (**G**) KEGG enrichment analysis of the downregulated genes in alveolar macrophages from COPD patients versus healthy controls.

Statistical analysis was performed in E and F using Mann-Whitney U test.

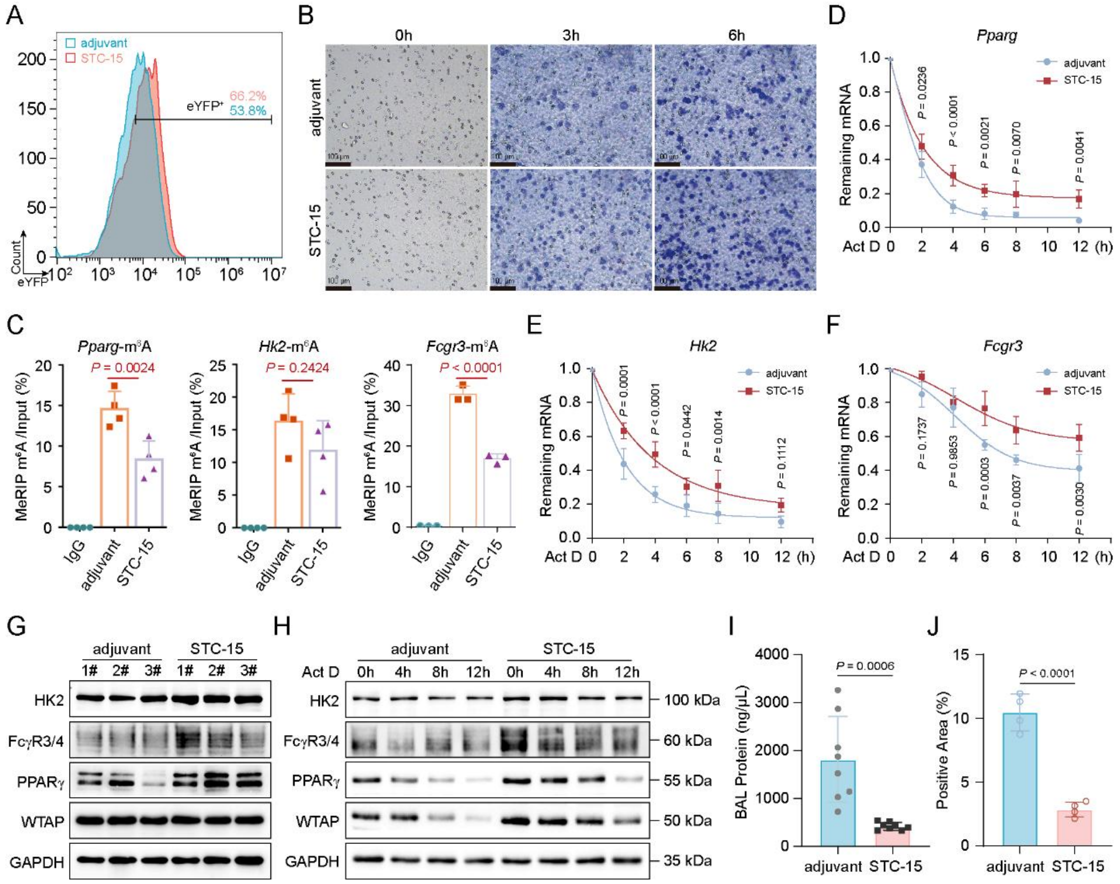


**Supplemental Figure 12. Inhibition of m$^6$A modification enhances antibacterial activity.** (**A**) Flow cytometry quantifying the phagocytic ability of adjuvant- or STC-15-treated AMs. (**B**) Representative imaging of migrated adjuvant- or STC-15-treated AMs after crystal violet staining. Scale bar, 100 μm. (**C**) MeRIP-qPCR showing the abundance of *Pparg*, *Hk2* and *Fcgr3* transcripts in adjuvant- or STC-15-treated AMs. (**D** to **F**) qRT–PCR showing the mRNA abundance of *Pparg*, *Hk2* and *Fcgr3* in adjuvant- or STC-15-treated AMs treated with Act D at the indicated time points. (**G**) Immunoblotting analysis of the expression of PPARγ, HK2 and FcγR3 in adjuvant- or STC-15-treated AMs. (**H**) Immunoblotting analysis of the expression of PPARγ, HK2 and FcγR3 in adjuvant- or STC-15-treated AMs treated with Act D at the indicated time points. (**I**) Quantification of BALF protein of control or STC-15 pre-treated mice infected with *P. aeruginosa*. n = 8 mice per group. (**J**) Quantification of MPO staining of the lung tissues from control or STC-15 pre-treated mice 12 hours after *P. aeruginosa* administration. n = 4 mice per group.

Data are presented as the mean ± s.d. in C, I and J with individual measurements overlaid as dots, and statistical analysis was performed in C, I and J using a two-tailed Student's *t*-test or performed in D-F using Multiple Comparisons Following Two-Way ANOVA. Data in A, B, G and H are representative of three independent biological experiments.

### Supplementary tables and table legends

**Supplementary Table 1. Information about reagents and resources**

| Reagent or Resource | Source | Identifier |
|---|---|---|
| **Antibodies** | | |
| WTAP | Proteintech | Cat# 10200-1-AP |
| EEA1 | Proteintech | Cat# 68065-1-Ig |
| HK2 | Proteintech | Cat# 22029-1-AP |
| PPARγ | Proteintech | Cat# 16643-1-AP |
| CD16/32 (FcγR3 + FcγR4) | Proteintech | Cat# 33284-1-AP |
| GAPDH | Proteintech | Cat# 10494-1-AP |
| Goat anti-Mouse IgG | Proteintech | Cat# SA00001-1 |
| Goat anti-Rabbit IgG | Proteintech | Cat# SA00001-2 |
| $m^6A$ antibody | CST | Cat# 56593 |
| FITC anti-mouse CD45 | Biolegend | Cat# 147709 |
| Ly6G-APC | Biolegend | Cat# 127614 |
| Cd11b-PE | BD | Cat# 557397 |
| Cd11b-FITC | Thermo | Cat# 11-0112-82 |
| Siglec F-PE | Thermo | Cat# 12-1702-82 |
| Cd11c-FITC | Thermo | Cat# 11-0114-82 |
| **Reagents** | | |
| LPS (From E. coli O111:B4) | InvivoGen | Cat# tlrl-eblps |
| 4% paraformaldehyde | Meilunbio | Cat# MA0192 |
| STC-15 | Selleck | Cat# E1728 |
| High Capacity Streptavidin Agarose Resin | Thermo Fisher | Cat# 20359 |
| Recombinant Murine GM-CSF | PeproTech | Cat# 315-03 |
| Actinomycin D | Sigma-Aldrich | Cat# A1410 |
| TRIzol reagent | Invitrogen | Cat# 15596026 |
| proteinase K | Roche | Cat# R011741 |
| pHrodo™ Red | ThermoFisher | Cat# P36600 |

| | | |
|---|---|---|
| PKH26 dye | Sigma–Aldrich | Cat# PKH26PCL |
| Hoechst 33342 | Beyotime | Cat# C1028 |
| **Experimental models: Organisms/strains** | | |
| C57BL/6 mice | Sun Yat-sen University Laboratory Animal Center | N/A |
| *LyzM*-Cre$^{+}$ *Wtap*$^{\Delta 1\text{-}77}$ mice (C57BL/6 background) | Generated for this study by Gempharmatech Co., Ltd.. | N/A |
| *Pseudomonas aeruginosa* | ATCC | ATCC27853 |
| *Escherichia coli* | ATCC | ATCC11303 |
| Influenza A virus (strain A/Puerto Rico/8/1934, H1N1) | Gift from Dr. Nan Qi | N/A |
| MDCK cells | ATCC | N/A |
| **Critical commercial assays** | | |
| *Evo M-MLV* RT Mix Kit with gDNA Clean for qPCR | AG | Cat# AG11728 |
| 2×Polarsignal® qPCR mix | MIKX | Cat# MKG802 |
| Magna MeRIP m$^6$A Kit | Merck Millipore | Cat# 17-10499 |
| RNeasy kit | QIAGEN | Cat# 74104 |
| Magic Red Cathepsin-B Assay Kit | Immunochemistry Tech | Cat# ICT-938 |
| Seahorse XF Cell Mito stress test kit | Agilent | Cat# 103015-100 |
| Seahorse XF glycolytic stress test kit | Agilent | Cat# 103020-100 |
| EpiQuik CUT&RUN m$^6$A RNA Enrichment (MeRIP) Kit | EPIGENTEK | Cat# P-9018 |

**Supplementary Table 2. List of oligonucleotides used in this study**

| Primer name | Primer sequence (5’→3’) |
| --- | --- |
| **qPCR primer** | |
| *Pparg* Forward | TCGCTGATGCACTGCCTATG |
| *Pparg* Reverse | GAGAGGTCCACAGAGCTGATT |
| *Fcgr3* Forward | CAGAATGCACACTCTGGAAGC |
| *Fcgr3* Reverse | GGGTCCCTTCGCACATCAG |
| *Hk1* Forward | AGGGCGCATTACTCCAGAG |
| *Hk1* Reverse | CCCTGTGGGTGTCTTGTGTG |
| *Hk2* Forward | TGATCGCCTGCTTATTCACGG |
| *Hk2* Reverse | AACCGCCTAGAAATCTCCAGA |
| *Hk3* Forward | CAGGGGACCTACAGGATTGAT |
| *Hk3* Reverse | GAGCATCTTCGTCATAGAAGGAG |
| *Pfkl* Forward | GAACTACGCACACTTGACCAT |
| *Pfkl* Reverse | CTCCAAAACAAAGGTCCTCTGG |
| *Pfkm* Forward | TGTGGTCCGAGTTGGTATCTT |
| *Pfkm* Reverse | GCACTTCCAATCACTGTGCC |
| *Pgm1* Forward | CAGAACCCTTTAACCTCTGAGTC |
| *Pgm1* Reverse | CGAGAAATCCCTGCTCCCATAG |
| *Tpi1* Forward | CCAGGAAGTTCTTCGTTGGGG |
| *Tpi1* Reverse | CAAAGTCGATGTAAGCGGTGG |
| *Pgk1* Forward | ATGTCGCTTTCCAACAAGCTG |
| *Pgk1* Reverse | GCTCCATTGTCCAAGCAGAAT |
| *Aldob* Forward | GAAACCGCCTGCAAAGGATAA |
| *Aldob* Reverse | GAGGGTCTCGTGGAAAAGGAT |
| *Eno1* Forward | TGCGTCCACTGGCATCTAC |
| *Eno1* Reverse | CAGAGCAGGCGCAATAGTTTTA |
| *Pdhb* Forward | GTGGAAGAAATACGGTGACAAGA |
| *Pdhb* Reverse | GGAGTAAAGGCAAACACCACAA |

| | |
|---|---|
| *Cs* Forward | GGACAATTTTCCAACCAATCTGC |
| *Cs* Reverse | TCGGTTCATTCCCTCTGCATA |
| *Idh3a* Forward | TGGGTGTCCAAGGTCTCTC |
| *Idh3a* Reverse | CTCCCACTGAATAGGTGCTTTG |
| *Ogdh* Reverse | GTTTCTTCAAACGTGGGGTTCT |
| *Ogdh* Forward | GCATGATTCCAGGGGTCTCAAA |
| *Sucla2* Forward | ACCCTTTCGCTGCATGAATAC |
| *Sucla2* Reverse | CCTGTGCCTTTATCACAACATCC |
| *Sdha* Forward | GGAACACTCCAAAAACAGACCT |
| *Sdha* Reverse | CCACCACTGGGTATTGAGTAGAA |
| *Sdhb* Forward | AATTTGCCATTTACCGATGGGA |
| *Sdhb* Reverse | AGCATCCAACACCATAGGTCC |
| *Sdhd* Forward | TGGTCAGACCCGCTTATGTG |
| *Sdhd* Reverse | GGTCCAGTGGAGAGATGCAG |
| *Cpt1a* Forward | CTCCGCCTGAGCCATGAAG |
| *Cpt1a* Reverse | CACCAGTGATGATGCCATTCT |
| *Acacb* Forward | CGCTCACCAACAGTAAGGTGG |
| *Acacb* Reverse | GCTTGGCAGGGAGTTCCTC |
| *Echs1* Forward | TTGTGAACTTGCCATGATGTGT |
| *Echs1* Reverse | TGCTCGGGTGAGTCTCTGAG |
| *Hahda* Forward | TGCATTTGCCGCAGCTTTAC |
| *Hadha* Reverse | GTTGGCCCAGATTTCGTTCA |
| *Hahdb* Forward | TGGTGGTGTTGAGTTAATGTCTG |
| *Hadhb* Reverse | TGTTCCATTCGAGAAACAGCAA |
| *Acaa2* Forward | CTGCTACGAGGTGTGTTCATC |
| *Acaa2* Reverse | AGCTCTGCATGACATTGCCC |
| **MeRIP–qPCR primer** | |
| *Pparg*-m$^6$A site Forward | ATCCGTAGAAGCCGTGCAA |
| *Pparg*-m$^6$A site Reverse | GCTCCATAAAGTCACCAAAGGG |

| | |
|---|---|
| *Hk2*-m$^6$A site Forward | ACAGCACACTGTCTCATGCT |
| *Hk2*-m$^6$A site Reverse | CTAGAGGGTAGCCACGGAGT |
| *Fcgr3*-m$^6$A site Forward | GGTGGAGGTCAGAGGACAAC |
| *Fcgr3*-m$^6$A site Reverse | GGAGGATCGCCAGTTCAAGG |
| *Fcgr4*-m$^6$A site Forward | GGGTCTCCATCCATGTTTCC |
| *Fcgr4*-m$^6$A site Reverse | GAGGCTCATGGGTTCTACAC |
